# PRDM16 controls smooth muscle cell fate in atherosclerosis

**DOI:** 10.1101/2025.02.19.639186

**Authors:** Josephine M.E. Tan, Lan Cheng, Ryan P. Calhoun, Angela H. Weller, Karima Drareni, Skylar Fong, Eirlys Barbara, Hee-Woong Lim, Chenyi Xue, Hanna Winter, Gaëlle Auguste, Clint L. Miller, Muredach P. Reilly, Lars Maegdefessel, Esther Lutgens, Patrick Seale

**Affiliations:** Institute for Diabetes, Obesity & Metabolism; Department of Cell and Developmental Biology; Perelman School of Medicine, University of Pennsylvania, Philadelphia, PA, USA; Department of Biomedical Informatics, Cincinnati Children’s Hospital Medical Center, Cincinnati, OH, USA; Department of Pediatrics, University of Cincinnati, Cincinnati, OH, USA; Division of Cardiology, Department of Medicine, Columbia University Irving Medical Center, Irving Institute for Clinical and Translational Research, Columbia University, New York, NY, USA; Institute of Molecular Vascular Medicine, University Hospital rechts der Isar, Technical University Munich, Germany; German Center for Cardiovascular Research (DZHK), partner site Munich Heart Alliance, Germany; Department of Genome Sciences; Department of Biochemistry and Molecular Genetics; University of Virginia, Charlottesville, VA 22908, USA; Experimental Cardiovascular Immunology Laboratory, Department of Cardiovascular Medicine and Immunology, Mayo Clinic, Rochester, MN, USA

## Abstract

Vascular smooth muscle cells (SMCs) normally exist in a contractile state but can undergo fate switching to produce various cell phenotypes in response to pathologic stimuli^1–3^. In atherosclerosis, these phenotypically modulated SMCs regulate plaque composition and influence the risk of major adverse cardiovascular events^4,5^. We found that PRDM16, a transcription factor that is genetically associated with cardiovascular disease, is highly expressed in arterial SMCs and downregulated during SMC fate switching in human and mouse atherosclerosis. Loss of *Prdm16* in SMCs of mice activates a synthetic modulation program under homeostatic conditions. Single cell analyses show that loss of *Prdm16* drives a synthetic program in all SMC populations. Upon exposure to atherogenic stimuli, SMC-selective *Prdm16* deficient mice develop SMC-rich, fibroproliferative plaques that contain few foam cells. Acute loss of *Prdm16* results in the formation of collagen-rich lesions with thick fibrous caps. Reciprocally, increasing PRDM16 expression in SMCs blocks synthetic processes, including migration, proliferation, and fibrosis. Mechanistically, PRDM16 binds to chromatin and decreases activating histone marks at synthetic genes. Altogether, these results define PRDM16 as a critical determinant of SMC identity and atherosclerotic lesion composition.

## Introduction

Cardiovascular diseases (CVDs) are leading causes of death and disability worldwide. The underlying cause of the majority of CVD is atherosclerosis, which refers to the build-up of fatty plaques in the artery walls^6,7^. Lesion development begins with the activation of endothelial cells in response to dyslipidemia and other pathologic stimuli, leading to monocyte adhesion, migration, and transformation into pro-inflammatory foam cells^8,9^. In this atherogenic milieu, arterial smooth muscle cells (SMCs), which normally exist in a quiescent or contractile state, undergo “phenotypic switching” or “modulation”, to produce various cell types. Synthetic modulated SMCs migrate, proliferate, and secrete extracellular matrix (ECM) components to help create a fibrous cap that surrounds lesions. SMCs can also give rise to “macrophage-like” and “osteogenic” cells in plaques^1–3^.

Modulated SMCs account for 40-70% of plaque cells and play a central role in regulating plaque composition and disease outcomes^4,5^. Rupture of an atherosclerotic plaque can produce a thrombus and cause a potentially lethal cardiovascular event such as myocardial infarction or stroke. Autopsy studies indicate that ∼70% of thrombi are caused by ruptured plaques, and that ∼80% of those plaques are thin-capped fibroatheromas, composed of a large lipid or necrotic core with immune infiltration, covered by a thin fibrous cap with decreased SMCs^10,11^. Thick fibrous caps rich in synthetic SMCs are thought to protect against rupture, whereas elevated immune cell activity and macrophage-like SMCs are associated with the development of vulnerable plaques^12,13^. The other main cause of atherothrombosis is plaque erosion, characterized by endothelial loss, low inflammation and high synthetic SMC content^11,14^. Here, the role of synthetic SMCs is less explored, and it remains unclear whether their increased number is protective or deleterious. Because of the significance of synthetic SMC modulation in vascular disease, identifying and understanding the mechanisms that regulate this phenotype switch is a critical area of current research.

Recent genome wide association studies (GWAS), combined with single-nucleus chromatin accessibility profiling of patient arteries, nominated *PR-domain containing 16* (*PRDM16*) as a candidate driver gene for coronary artery disease, potentially acting in SMCs^15^. PRDM16 is an epigenetic and transcriptional regulator that has been extensively studied in adipocytes, where it drives an energy-burning metabolic program^16–18^. PRDM16 also regulates mitochondrial and metabolic programs in other cell types, including cardiomyocytes, intestinal epithelial cells, and hematopoietic stem cells^19–22^. Recently, PRDM16 has also been implicated in regulating ventricular cardiomyocyte and arteriovenous endothelial cell (EC) function^23,24^.

Here, we investigated the role of PRDM16 in regulating SMC identity and atherosclerotic lesion formation. We found that PRDM16 is highly expressed in arterial SMCs and downregulated during atherosclerosis-induced SMC modulation in mice and humans. Acute or chronic loss of PRDM16 in SMCs of mice activated synthetic process-related genes in SMCs under homeostatic (basal) conditions. Under atherosclerotic conditions, these *Prdm16* deficient animals developed fibrous lesions, characterized by the increased proliferation of synthetic modulated SMCs, extensive collagen deposition and reduced foam cell formation. Single cell analyses revealed that the loss of *Prdm16* upregulated synthetic genes across all SMC populations and increased the formation of “synthetic modulated cells” during atherosclerosis. Conversely, increased expression of PRDM16 in cell models powerfully blocks the fibrotic response through direct chromatin binding to synthetic genes. Overall, these results define PRDM16 as a critical repressor of the synthetic phenotype in SMCs and a determinant of atherosclerotic plaque composition.

## Results

### *PRDM16* is a cardiovascular disease-associated gene with enriched expression in SMCs

GWAS have identified several single nucleotide polymorphisms (SNPs) in the *PRDM16* locus that are associated with CVD risk (Supplemental Fig. 1a). Moreover, rare *PRDM16* coding variants identified in the UK Biobank are associated with atherosclerotic cardiovascular diseases, e.g. coronary artery disease, stroke, and cerebrovascular disease. Analysis of *PRDM16* mRNA expression levels in human tissues using the GTEx portal revealed conspicuously high levels in arteries, including aorta, coronary arteries, and tibial arteries, as compared to other tissues (Fig. 1a). Single cell expression analysis of human arteries showed a striking (>10-fold) enrichment of *PRDM16* mRNA levels in SMCs relative to endothelial cells or fibroblasts (Fig. 1b).

**Figure 1.**
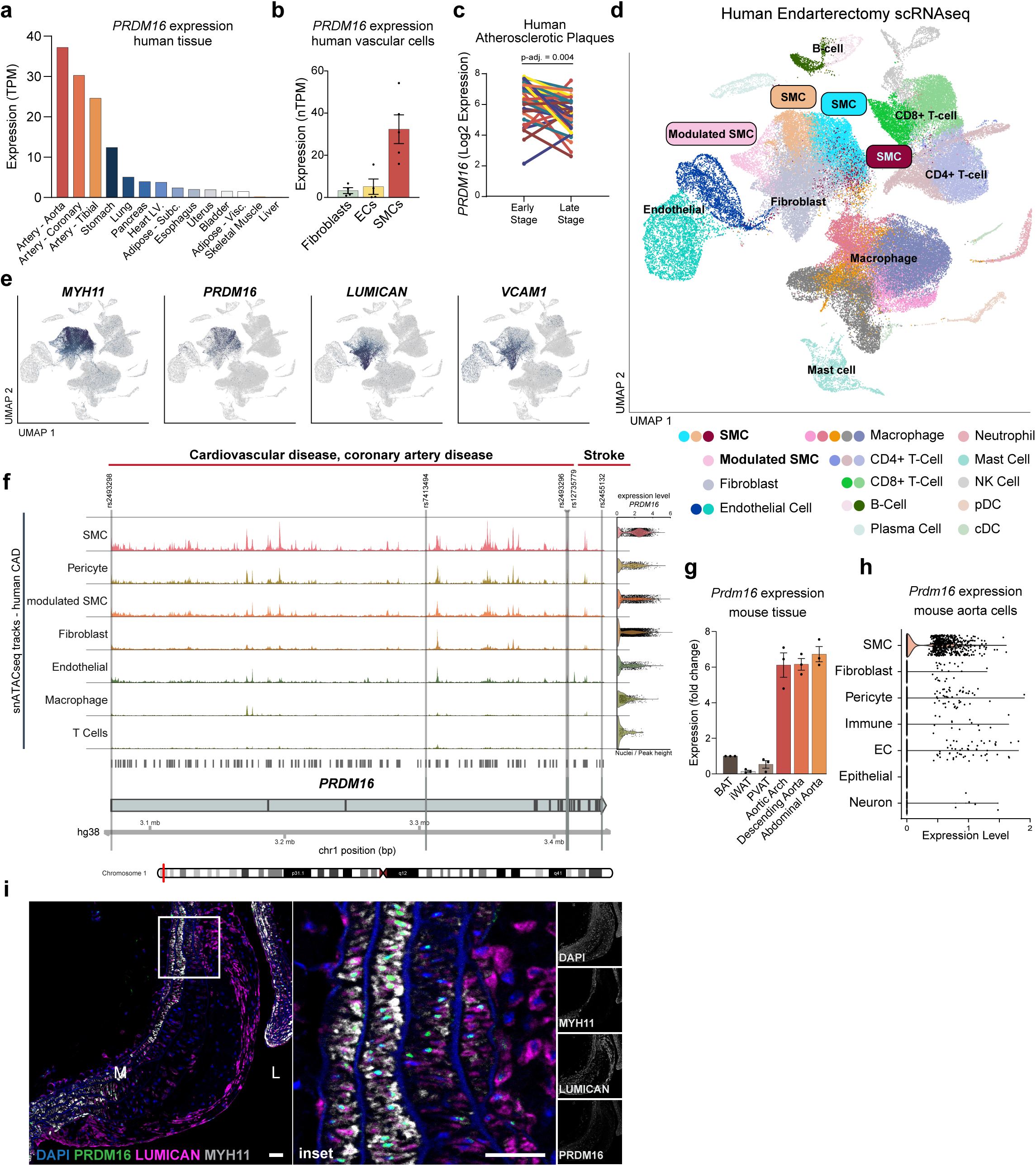
PRDM16 expression decreases during SMC modulation in mouse and human atherosclerosis. **a,** *PRDM16* expression in whole human tissues from The Genotype-Tissue Expression (GTEx) database in transcripts per million (TPM) (dbGaP Accession phs000424.v8.p2). **b,** Expression of *PRDM16* in vascular cell types in number of transcripts per million (nTPM) per cluster (Human Protein Atlas). Fibroblasts (*PDGFRA, PI16*+), ECs (*PECAM1, BMP4*+), SMCs (*MYH11, CNN1*+). **c,** Expression of *PRDM16* in endarterectomy samples of early and late-stage lesions of the same human patients 25, n= 38. **d,** UMAP visualization of clusters identified by scRNAseq of human endarterectomy samples 26 n=21. **e**, UMAP feature plots showing expression of SMC markers *MYH11, PRDM16,* and modulated SMC markers *LUMICAN* and *VCAM1.* **f,** Visualization of snATACseq tracks from human coronary artery samples (N=41 patients)15, clustered by cell-type at the *PRDM16* locus, showing the locations of SNPs associated with and/or CVD, and stroke are indicated. **g,** RT-qPCR analysis of *Prdm16* mRNA expression in mouse brown adipose tissue (BAT), inguinal white adipose tissue (iWAT), perivascular adipose tissue (PVAT), aortic arch, descending aorta, and abdominal aorta. N=3 mice. Data is presented as mean ± S.E.M. **h,** Violin plot showing *Prdm16* expression level of indicated genes in cell clusters (TPM) identified from scRNAseq of mouse aortas 29. Smooth muscle cell (SMC), endothelial cell (EC). **i,** Representative sagittal section of aorta containing a lesion with a fibrous cap from mice treated with AAV8-PCSK9-D377Y and western diet for 12 weeks. Immunostaining for DAPI (blue), MYH11 (white), Lumican (magenta) and PRDM16 (green). Scale bar: 50 μm. M: tunica media, L: lumen.

### PRDM16 is downregulated during SMC phenotypic modulation in atherosclerosis

SMCs undergo phenotypic modulation during atherosclerosis. This process involves the downregulation of canonical SMC marker genes such as *MYH11, TAGLN* and *CNN1* and activation of diverse gene programs that promote vascular remodeling, plaque formation, and fibrous cap formation. Paired analysis of early-versus late-stage atherosclerotic lesions from patients^25^ showed lower *PRDM16* levels in late-stage lesions, which contain a higher proportion of modulated SMCs (Fig. 1c). Evaluation of human plaques by single cell expression profiling^26^ showed that *PRDM16* was highly expressed in canonical SMCs (*MYH11*+, *TAGLN*+), and decreased in modulated SMCs (Lumican, *LUM*+, *COL1A1*+) (Fig. 1d-e). Analysis of a second scRNAseq dataset from human carotid artery plaques^27^ showed high *PRDM16* levels in contractile SMCs, and greatly diminished expression levels in the two modulated SMC clusters, “modulated” and “fibroblast-like modulated SMCs” (Supplemental Fig. 1b-d). Single nucleus transposase-accessible chromatin (ATAC)seq analysis of CAD patient samples^15^ showed the highest levels of chromatin accessibility at the *PRDM16* locus in SMCs, with lower accessibility in pericytes and modulated SMCs (Fig. 1f). Significantly, multiple CVD-associated SNPs in *PRDM16* localized to accessible chromatin regions in SMCs (Supplemental Fig. Fig. 1a, Fig. 1f).

The *Prdm16* expression pattern was similar in mice, with higher levels of *Prdm16* mRNA detected in all regions of the aorta compared to tissues where it has been most studied, such as interscapular brown adipose tissue (BAT), perivascular adipose tissue (PVAT) and subcutaneous white adipose tissue (WAT) (Fig. 1g). Single-cell analysis of mouse aortae^28^ showed that *Prdm16* expression was highly enriched in SMCs relative to other vascular cell types (Fig. 1h, Supplemental Fig. 1e-f). We then assessed PRDM16 expression in mouse atherosclerosis. Immunofluorescence (IF) staining of lesions from two different mouse models of atherosclerosis showed that modulated SMCs, marked by Lumican expression, were concentrated in the fibrous cap, well segregated from MYH11+ SMCs in the media (Fig. 1i; Supplemental Fig. 1g). PRDM16 protein was localized in the nucleus of MYH11+ cells in the media and lost from Lumican+ modulated SMCs in lesions. Interestingly, PRDM16 protein levels decreased across the width of lesions from high levels in vessel wall SMCs to undetectable expression in the distal fibrous cap cells (Fig. 1i; Supplemental Fig. 1g). In conclusion, PRDM16 is highly expressed in arterial SMCs and downregulated during SMC modulation and fibrotic cap formation in human and mouse atherosclerosis.

### PRDM16 represses the synthetic gene program in SMCs under homeostatic conditions

To study the role of PRDM16 in SMCs, we generated mice with SMC-selective constitutive deletion of *Prdm16* using *Tagln* Cre (SMC*-*cKO). IF analysis of aortae from these mice showed efficient ablation of PRDM16 in SMCs, while endothelial cell expression of PRDM16 remained intact (Supplemental Fig. 2a). SMC-cKO mice were born at expected Mendelian ratios, appeared healthy, maintained the same bodyweight as control littermates and displayed normal glycemic control, as determined by glucose tolerance testing (Supplemental Fig. 2b-c). Histological examination of aortae showed a ∼30% reduction in medial thickness (Supplemental Fig. 2d-e), and decreased Elastin staining in SMC-cKO compared to control mice (Supplemental Fig. 2f). Systolic blood pressure was slightly lower in SMC-cKO compared to control mice but remained within the normal range (90-120 mmHg) (Supplemental Fig. 2g).

To determine how PRDM16 affects SMC gene expression, we performed RNA sequencing on cleaned aortae (stripped of perivascular tissue and endothelium), from control and SMC-cKO mice. Canonical SMC marker genes such as *Myh11* and *Tagln* were amongst the highest expressed genes and were expressed at similar levels in control and SMC-cKO mice (Fig. 2a). Many genes related to myofibroblast transition, fibrosis and ECM remodeling (e.g. *Ankrd1, Col12a1, Mustn1, Chrdl1, Des, Coro6 and Egr1*) were upregulated in SMC-cKO aortae (Fig. 2a, Supplemental Fig. 2h). Gene ontology (GO) analysis of the upregulated genes in cKO aortae identified processes that converge on synthetic SMC modulation, such as “cell migration”, “regulation of cell population proliferation”, “response to wounding” and “extracellular matrix organization” (Fig. 2b). Gene set enrichment analysis of the upregulated genes identified “Epithelial Mesenchymal Transition”, a process related to synthetic SMC switching, as a top hallmark pathway (Fig. 2c). SMC-cKO aortae also expressed elevated levels of some genes involved in cardiac/circulatory system development and markers of secondary heart field-derived SMCs, such as *Tnnt2* and *Myo18b* (Fig. 2a, b). The top GO pathways for the genes downregulated in cKO aortae corresponded to nonspecific transcriptional and sensory/perception processes (Supplemental Fig. 2h). *Prdm16* deficiency did not affect the expression of either *Tcf21* or *Klf4,* which are known transcriptional regulators of SMC modulation^4,29^ (Supplemental Fig. 2i). Additionally, genes related to the “macrophage-like SMC switch” (e.g. *Lgals3*, *Cd68, Vcam1*) were unaffected by *Prdm16*-deficiency (Supplemental Fig. 2i).

**Figure 2.**
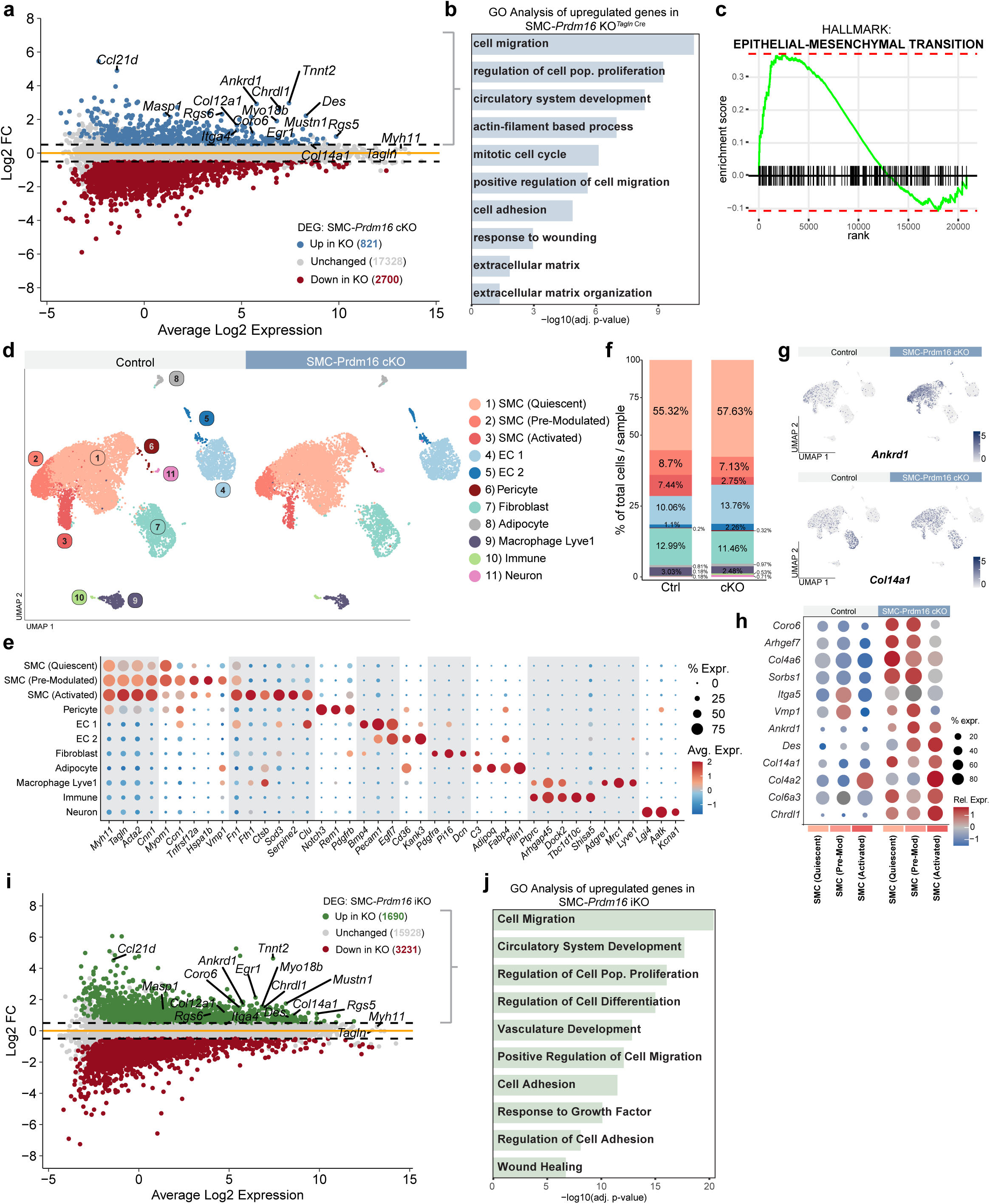
PRDM16 represses the synthetic gene program in SMCs under homeostatic conditions. **a,** Expression MA plot of genes associated with synthetic SMC processes and muscle identity in aortas from control and and cKO mice (n=5 per group). Genes upregulated in cKO mice are in blue (log2FC > 0.5 & padj. < 0.1), downregulated genes in cKO are red (log2FC < −0.5 & padj. < 0.1), grey is unchanged. **b,** (Gene ontology (GO) analyses of genes upregulated in and cKO vs. control aortas. N=5 mice per group. **f,** Gene set enrichment analysis showing gene enrichment for hallmark process epithelial to mesenchymal transition in cKO vs. control aortas. **d,** UMAP visualization of scRNAseq expression data from whole aorta of control (n=4, 11,297 cells) and cKO mice (n=4, 9,118 cells), revealing 11 clusters. **e,** Dot plot displaying the relative expression of cluster-defining marker genes across identified clusters. Identified clusters are shown on the y-axis and marker genes on the x-axis. Dot size represents the proportion of cells in a cluster expressing each gene, and a color scale from red (high) to blue (low) indicates relative expression levels. Legends for relative expression and cell proportions appear on the right. **f,** Bar charts representing the proportion of cells assigned to each cluster relative to the total number of cells analyzed under each condition. Clusters are color-coded and labelled according to (d). **g,** UMAP feature plot of *Ankrd1* and *Col14a1* expression in clusters of (d) in control vs cKO aortae. **h,** Dot plot displaying synthetic genes across SMC clusters in control and cKO conditions. Expression levels and cell proportions are plotted for each SMC cluster: quiescent, pre-modulated, and activated. Dot size represents the proportion of cells expressing each gene within a cluster, while a color scale indicates relative expression levels. **i,** Expression MA plot of genesassociated with synthetic SMC processes and muscle identity in aortas from control and and iKO mice (n=3 per group). Genes upregulated in iKO mice are in green (log2FC > 0.5 & padj. < 0.1), downregulated genes in iKO are red (log2FC < −0.5 & padj. < 0.1), grey is unchanged. **j,** (Gene ontology (GO) analyses of genes upregulated in and iKO vs. control aortas. N=3 mice per group.

We next utilized the 10x Flex platform to perform single-cell RNA sequencing (scRNA-seq) on fixed aortae (spanning from the root to the iliac bifurcation) of control and cKO mice. A main advantage of this technique is that it preserves the transcriptional profiles of aortic cells in their native tissue context prior to cell dissociation. Unsupervised clustering of 20, 415 cells identified 7 major and 4 minor clusters (each <1% of cells per condition). The major clusters comprised 3 SMC subtypes, 2 EC populations, adventitial fibroblasts and a macrophage cluster marked by expression of the vascular-resident macrophage marker *Lyve1* (Fig. 2d-f). The largest SMC cluster corresponded to “Quiescent” SMCs, marked by the expression of canonical SMC marker genes (e.g. *Myh11*, *Tagln*, *Cnn1*) and *Myom1*. A second SMC cluster, which we termed “Pre-Modulated”, expressed canonical SMC markers, *Myom1*, along with elevated levels of synthetic cell marker genes (e.g. *Ccn1*, *Tnfrsf12a, Vmp1* and *Fn1*). The third SMC cluster, which we termed “Activated”, expressed high levels of canonical SMC markers (e.g. *Myh11*, *Cnn1*), some synthetic genes (e.g. *Fn1*, *Col4a2*), and certain genes associated with ECM remodeling (e.g. *Ctsb*, *Clu*, and *Sod3*) (Fig. 2e, Supplemental Fig. 2j). We designed custom probes to detect the floxed exon of *Prdm16* (exon 9), which showed a specific loss of *Prdm16* expression from all three SMC clusters (Supplemental Fig. 2j). Most strikingly, synthetic genes (e.g. *Ankrd1, Col14a1*, *Col4a6, Coro6, Tnnt2, Chrdl1*) that were elevated in whole *Prdm16* cKO aortae were markedly and broadly upregulated across all 3 SMC clusters of cKO versus control mice (Fig. 2g,h, Supplemental Fig. 2j). While pre-modulated and activated SMCs from control mice expressed detectable levels of certain synthetic cell marker genes, the levels of these genes were further upregulated by *Prdm16*-deficiency (Fig. 2g,h, Supplemental Fig. 2j). Thus, the loss of *Prdm16* promotes the expression of synthetic genes across all types of SMCs under homeostatic conditions.

To evaluate the effects of acute *Prdm16*-deletion in SMCs of adult animals, we generated a tamoxifen-inducible *Myh11-Cre^ERT^*^2^-driven *Prdm16* KO mouse model (SMC-iKO). Adult (8-week-old) SMC-iKO and control mice were treated with tamoxifen to delete *Prdm16* specifically in SMCs. IF staining of aortae one week after tamoxifen treatment showed a near complete loss of PRDM16 protein expression specifically in SMCs of iKO mice (Supplemental Fig. 2k). Blood pressure, medial thickness and the elastic lamina were unchanged between control and iKO mice (Supplemental Fig. 2l-o). RNAseq analysis of cleaned aortae from control and SMC-iKO mice 1-week after tamoxifen treatment showed that acute loss of *Prdm16* upregulated a similar profile of genes to that observed in cKO mice, including synthetic genes (e.g. *Ankrd1, Coro6, Chrdl1, Col14a1*), *Tnnt2* and *Myo18b* (Fig. 2i, Supplemental Fig. 2p). GO analysis of the upregulated genes identified synthetic SMC modulation processes (e.g. migration, proliferation, adhesion, wound healing) (Fig. 2j). Altogether, these results indicate that PRDM16 represses a synthetic gene program in SMCs, suggesting a key regulatory role in SMC phenotype modulation.

### Loss of PRDM16 drives the development of SMC-rich and fibrous atherosclerotic plaques

We next sought to determine how the loss of PRDM16 in SMCs would impact atherosclerotic lesion development. We induced atherosclerosis in control and SMC-cKO mice by injection of AAV8-*mPCSK9* D377Y coupled with Western diet feeding under thermoneutral housing conditions (30°C) for 12 weeks. Thermoneutral housing provides a more humanized physiologic state by exempting mice from thermal stress, which lowers brown fat activity and decreases metabolic rate. Control and SMC-cKO mice gained weight at a similar rate during Western diet feeding (Supplemental Fig. 3a). Total plasma cholesterol and other plasma lipids (HDL, non-HDL, and triglycerides) were increased to equivalently high levels in control and KO mice (Supplemental Fig. 3b-e).

We assessed lesion development by analyzing serial sagittal sections of aortic arches from control and SMC*-*cKO mice (Supplemental Fig. 3f). Atherosclerotic plaques were readily detected in both mouse groups, and quantification of lesion area revealed no significant differences between control and *Prdm16* KO mice, though variability was higher in the KO group (Fig. 3a). H&E staining revealed striking morphological differences between control and KO lesions (Fig. 3b). Lesions from control mice displayed a classical fatty-streak morphology, with some showing a thin fibrous cap but generally not showing calcification or necrotic cores at this stage. By contrast, lesions from SMC*-*cKO mice were extremely dense, disorganized, and lacked foamy (lipid containing) cells. Indeed, in many cases, KO lesions were only distinguishable from the media by their disorganized fibrous structure and by their protuberance into the lumen (Supplemental Fig. 3g). Sirius red staining showed that control lesions contained low levels of collagen, which, when present, was confined to the fibrous cap (Fig. 3c,d). By contrast, KO lesions contained remarkably high amounts of collagen that was widely distributed (55% compared to 15% of lesion area) (Fig. 3c,d). IF staining for smooth muscle actin (SMA) showed that KO plaques contained much higher numbers and a wider distribution of modulated SMCs compared to control plaques (Fig. 3e, Supplemental Fig. 3h). Conversely, KO plaques contained fewer cells with lipid droplets (i.e. foamy cells), marked by expression of the lipid droplet protein Perilipin 2 (PLIN2) (Fig. 3e,f). We also assessed cell proliferation as a hallmark feature of SMC modulation. Control lesions contained small numbers of proliferating SMA+ cells, marked by Ki67 expression, that were confined to the junctional region between the media and lesion (Fig. 3g, Supplemental Fig. 3h). KO lesions contained ∼5-fold more proliferating SMA+ cells that were distributed throughout the lesion (Fig. 3g, Supplemental Fig. 3h).

**Figure 3.**
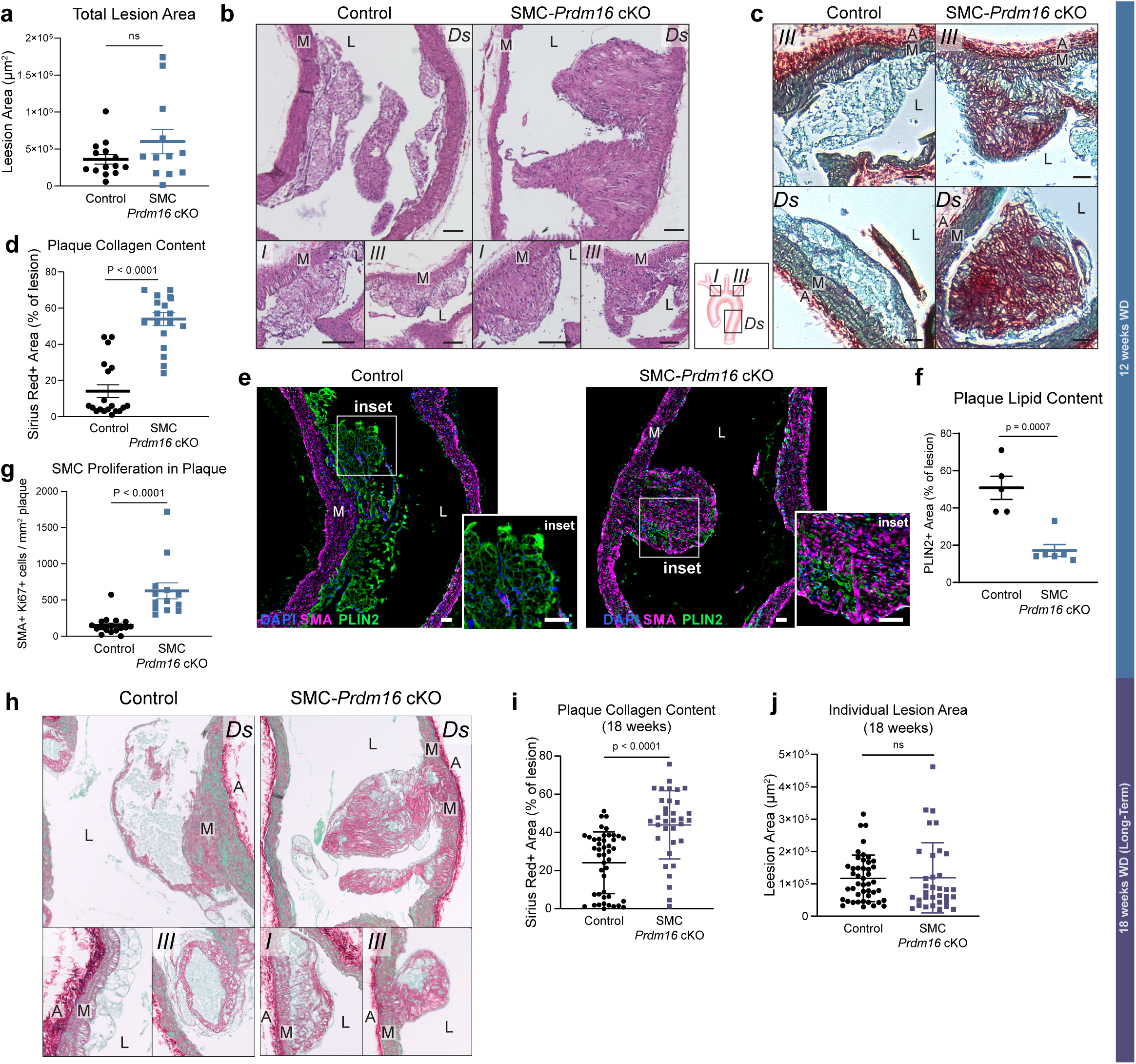
Loss of PRDM16 drives synthetic SMC modulation and fibrosis during moderate and advanced atherogenesis. **a,** Quantification of total lesion area of control and cKO mice injected with AAV8-*mPCSK9* D377Y after 12 weeks of Western Diet. This was measured by averaging the lesion area of 5 sequential sagittal sections (24 μm apart) per aorta (n = 14/12 Ctrl/KO). Data is represented as mean ± S.D. ns = not significant. **b,** H&E-stained sagittal sections of the aortic arch showing representative atherosclerotic lesions in control and KO mice at different locations. Images are representative of n = 14/12 Ctrl/KO. Scale bar: 100 μm. M: tunica media, L: lumen. A diagram of the aortic arch shows the orientation: I= brachiocephalic artery, III = left subclavian artery, Ds = descending aorta. **c,** Sagittal aortic arch sections stained with Sirius red/Fast-Green shows collagen deposition in representative lesions from control and KO mice. Scale bar: 50 μm. M: tunica media, L: lumen. **d,** Quantification shows the percentage of Sirius red positive area per lesion. Staining and quantification were performed on 5 mice per group, each dot represents a lesion. Data is represented as mean ± S.E.M. **e,** Immunostaining for Perilipin 2 (PLIN2) (green), smooth muscle actin/Acta2 (SMA) (magenta) and Dapi (blue, nuclei) in lesions from Control and KO mice. Representative of n=3 per group. Scale bar: 50 μm. M: tunica media, L: lumen. **f,** Quantification of PLIN2 positive area per lesion area. Each dot represents a lesion. **g,** Quantification of Ki67 and SMA double positive cells per lesion area. Each dot represents a lesion from n=5 mice per group. Data is represented as mean ± S.E.M. **h,** Sagittal aortic arch sections of control / cKO mice injected with AAV8-*mPCSK9* D377Y after 18 weeks of Western Diet. Sections were stained with Sirius red/Fast-Green and show collagen deposition in representative lesions from control and KO mice (n = 7/5). Scale bar: 50 μm. M: tunica media, L: lumen. **i,** Quantification shows the percentage of Sirius red positive area per lesion. Staining and quantification were performed on 7 control and 5 cKO mice, each dot represents a lesion. Data is represented as mean ± S.E.M. **d,** Individual size of each lesion identified in control and cKO mice (n = 5 and 7 respectively). each dot represents a lesion.

We further evaluated the effects of *Prdm16* deficiency at a more advanced stage of atherosclerosis (18 weeks). Control and cKO mice maintained similar body weights throughout the 18 weeks of Western diet feeding, and total lesion area was not changed (Supplemental Fig. 3i-k). Control plaques at 18 weeks displayed a more complex structure compared to those at 12 weeks, featuring a better defined fibrous cap, and an increase in collagen content (Fig. 3h,i). cKO lesions at 18 weeks still exhibited a significant increase in fibrosis (∼2-fold increase in Sirius red-positive area) compared to controls, but also contained more foam cells than cKO lesions at 12 weeks, reflecting more advanced lesion development and immune cell infiltration in long-term atherosclerosis (Fig. 3h,i). No change in the size of individual lesions was observed between control and long-term cKO mice, implying that the elevated proliferation in cKO SMCs mice did not lead to continuous lesion enlargement (Fig. 3j). Altogether, these results show that *Prdm16* deficiency promotes the formation of fibroproliferative, SMC-rich lesions with reduced foam cells.

### Inducible loss of *Prdm16* in adult mice promotes the formation of fibrotic plaques with thicker caps

We induced atherosclerosis in control and SMC-iKO mice 1-week following tamoxifen treatment via injection of AAV8-*mPCSK9* D377Y and Western diet feeding for 12 weeks. Body weights were similar between the groups until 12 weeks, when iKO mice weighed slightly less than controls (Supplemental Fig. 4a). Lesion area was unchanged between control and iKO mice (Fig. 4a,b). H&E staining showed that iKO lesions contained fewer foam cells with smaller lipid droplets, as compared to control lesions (Supplemental Fig. 4b). Notably, iKO lesions developed a prominent fibrous cap, whereas control lesions either entirely lacked a cap or possessed a thin cap (Fig. 4c, Supplemental Fig. 4b). Sirius red staining showed that iKO lesions contained much higher collagen levels than controls, with higher collagen deposition present throughout the lesion and in the fibrous caps (Fig. 4c,d). IF analysis showed elevated SMC content in iKO plaques accompanied by a reduction in PLIN2+ lipid-containing cells (Fig 4e,f).

**Figure 4.**
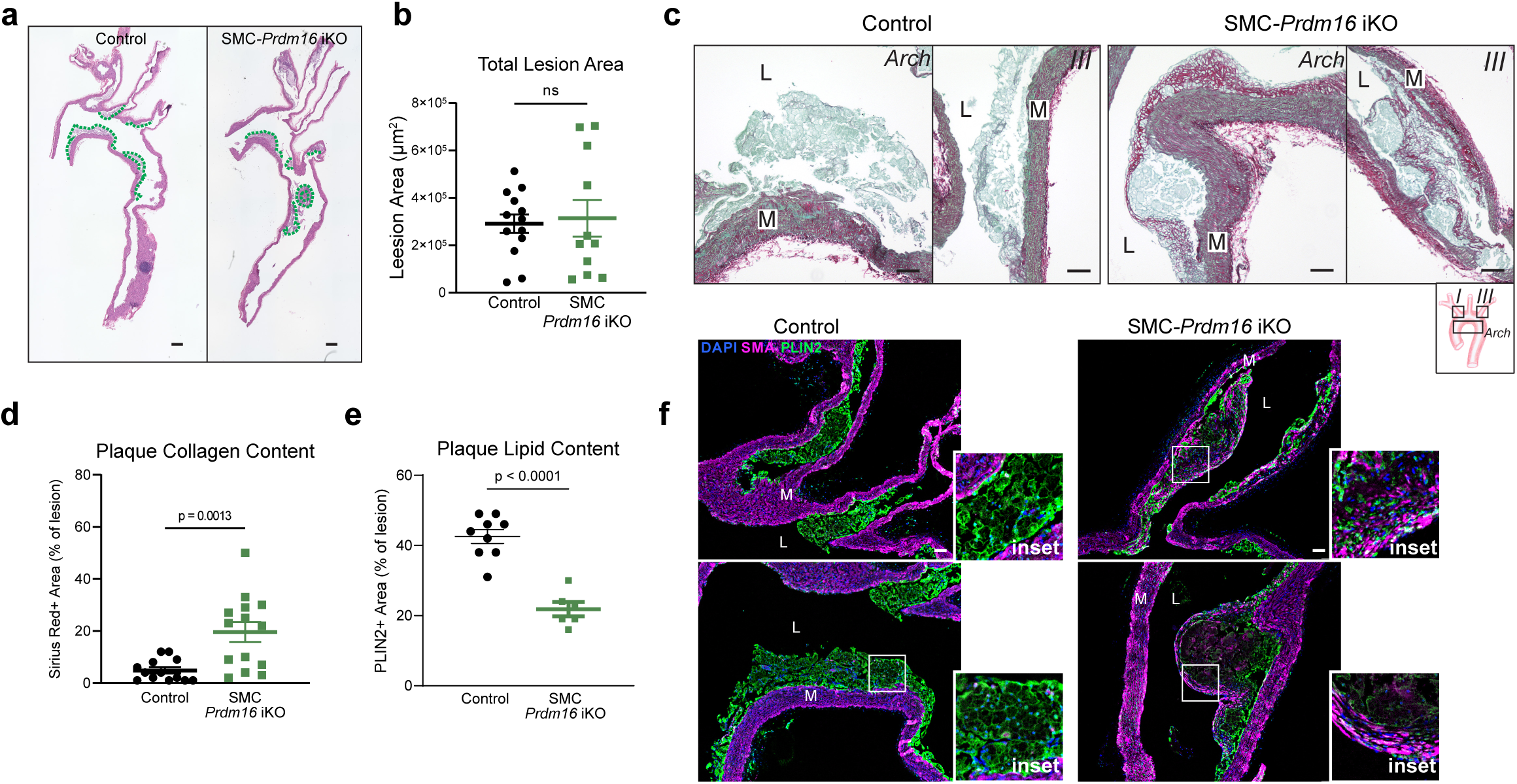
Acute loss of PRDM16 promotes synthetic SMC modulation and fibrosis. **a,** H&E staining of sagittal sections of the aortic arch from control and SMC-iKO mice at 12 weeks of HFD feeding. Lesion area is outlined in green. Only lesions in the arch, the start of the descending aorta and the beginnings of the branches were included. Scale bar: 100 μm (n = 13/11 Ctrl/iKO). **b,** Quantification of total lesion area, as measured by averaging the lesion area of 5 sequential sagittal sections (24 μm apart) per aorta (n = 13/11 Ctrl/KO). Data is presented as mean ± S.D. **c,** Sirius red/Fast-Green staining of aortic arch sections shows collagen depositions within lesions in control and iKO mice. Scale bar: 50 μm **d,** Quantification shows the percentage of Sirius red positive area per lesion. Staining and quantification were performed on sections 5 mice per group, each dot represents a lesion. Data is represented as mean ± S.E.M. **e,** Quantification of PLIN2 positive area per lesion area in control/iKO mice. Each dot represents a lesion. **f,** Immunostaining for Perilipin 2 (PLIN2) (green), smooth muscle actin/Acta2 (SMA) (magenta) and Dapi (blue, nuclei) in lesions from Control and iKO mice. Representative of n=3 per group. Scale bar: 50 μm. M: tunica media, L: lumen.

### PRDM16 represses SMC synthetic processes and fibrosis

*PRDM16* is highly expressed in mouse and human arterial SMCs *in vivo*, but negligibly expressed in cultured human coronary artery SMCs (hCaSMC) (Ct >35), providing a platform for gain-of-function assays (Supplemental Fig. 5a). A critical feature of synthetic SMC modulation during atherosclerosis is the migration of SMCs out of the media to form a fibrous cap. We tested the role of PRDM16 in SMC migration using a wound healing (scratch) assay. hCaSMCs were transduced with a control guide or a guide targeting the *PRDM16* promoter to activate its expression using CRISPR-activation (CRISPRa). *PRDM16* levels increased from near undetectable (Ct >35) to Ct ∼25, which is similar to *Prdm16* expression levels in mouse aortae (Supplemental Fig. 5a, d). Cells were grown to confluence, followed by inducing a wound and imaging migration with time-lapse microscopy. *PRDM16* activation significantly reduced SMC wound closure rate by about 20% (Fig. 5a, Supplemental Fig. 5b). We also tested if PRDM16 regulates proliferation and/or fibrosis, other synthetic processes involved in SMC modulation. *PRDM16*-CRISPRa cells displayed a small but significant decrease in proliferation compared to control cells (Fig. 5b). *PRDM16* CRISPRa cells also expressed lower levels of many synthetic/fibrosis genes (that were also upregulated in *Prdm16-*KO mouse aortae), including *COL16A1*, *TNFRSF11B*, *STARD13* and *ANKRD1* (Fig. 5c). Additionally, we used a lentiviral expression vector to attain higher *Prdm16* levels in hCaSMCs, which led to stronger decreases in cell migration (50%), proliferation (70%), and synthetic/fibrosis gene expression (Fig. 6d-f, Supplemental Fig. 5a,c,e). These results indicate that PRDM16 dose-dependently inhibits synthetic processes in SMCs.

**Figure 5.**
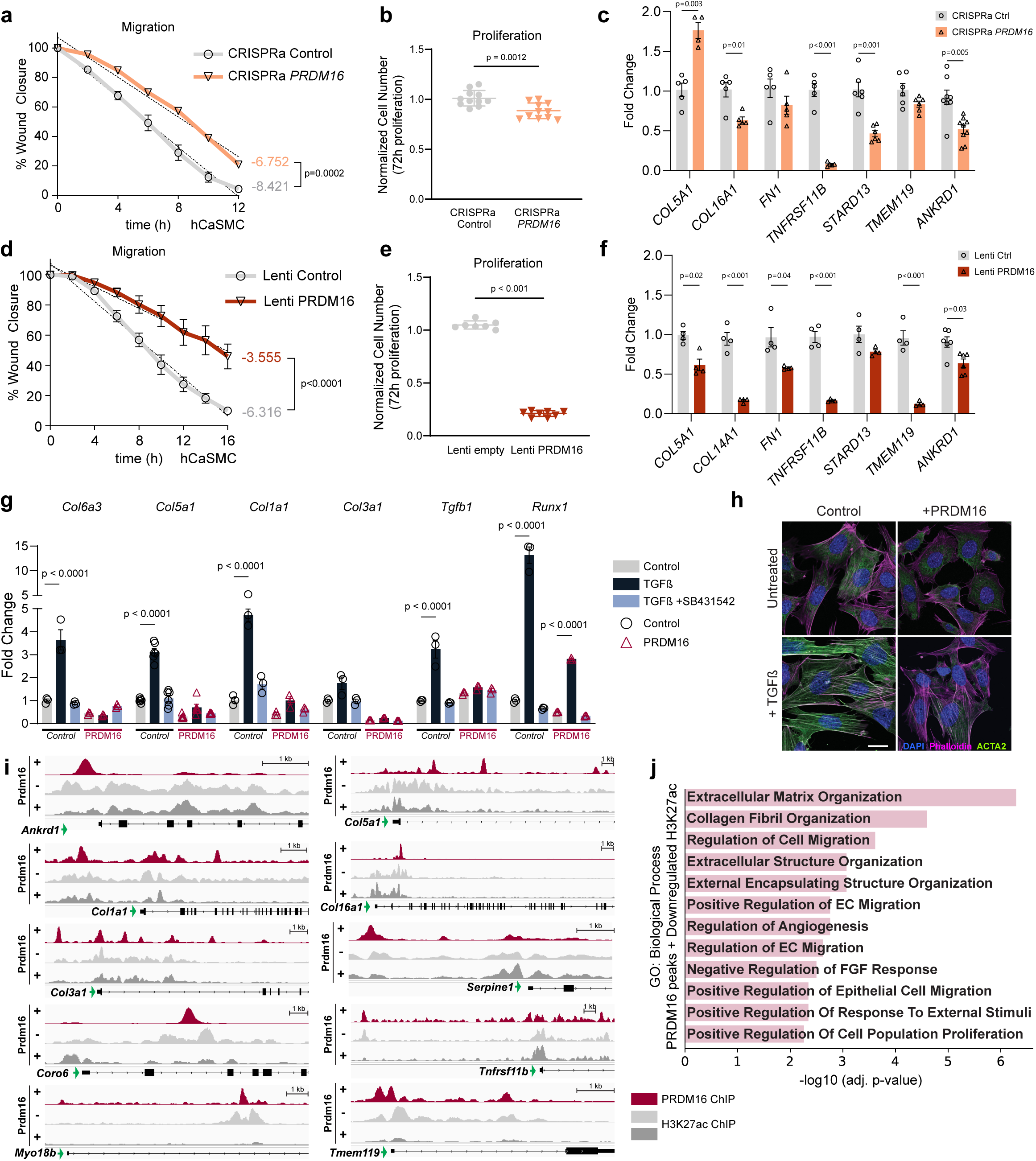
PRDM16 expression represses SMC synthetic processes and fibrosis. **a,** a migration assay using hCaSMCs transduced with a control CRISPRa plasmid or a vector expressing a CRISPRa guide for *PRDM16.* The cells were pre-incubated with the proliferation inhibitor mitomycin-C, a scratch was created and the cells were monitored using widefield microscopy for 12 hours. Scratch closure was quantified every 2 hours and the line graph represents the average percentage of wound closure after scratch. N = 6. Data is presented as mean ± S.E.M **b,** MTT assay of cell proliferation for hCaSMCs transduced with control or PRDM16 CRISPRa-expressing virus. N = 11. Data is presented as mean ± S.E.M. **c,** mRNA levels of indicated genes in control and PRDM16-CRISPRa-expressing hCaSMCs. **d,** a migration assay using hCaSMCs transduced with a control lentivirus or a lentivirus encoring HA-PRDM16. The cells were pre-incubated with the proliferation inhibitor mitomycin-C, a scratch was created and the cells were monitored using widefield microscopy for 16 hours. Scratch closure was quantified every 2 hours and the line graph represents the average percentage of wound closure after scratch. N = 5. Data is presented as mean ± S.E.M. **e,** MTT assay of cell proliferation for hCaSMCs transduced with the constructs used in d. N = 7/8 for control/lenti PRDM16. Data is presented as mean ± S.E.M. **f,** mRNA levels of indicated genes in control/lenti PRDM16-expressing hCaSMCs. **g,** mRNA levels of indicated genes in control and PRDM16-expressing fibroblasts treated with TGFß1 and/or SB4318542 or vehicle control for 48h. N > 9. Data is presented as mean ± S.E.M. **h,** Immunostaining for Phalloidin (magenta), ACTA2 (green), and DAPI (nuclei, blue) in control and PRDM16 expressing cells +/- recombinant TGFß1 for 48h. Scale bar: 10 μm. **i,** ChIP-seq tracks for PRDM16 and H3K27Ac at indicated synthetic genes in control and PRDM16-expressing fibroblasts. **j,** GO analysis of genes containing PRDM16 ChIP-seq peaks that display a concomitant decrease of H3K27ac in PRDM16-expressing vs. control cells.

We next investigated the effects of ectopic *Prdm16* expression in fibroblasts, which exhibit a highly synthetic phenotype. PRDM16 powerfully repressed the expression of synthetic genes, and other genes that were also repressed by PRDM16 in aortae (from RNAseq studies). To further stimulate fibrosis, we treated these cells with the canonical fibrosis inducer TGFß1. Treatment with TGFß1 markedly upregulated synthetic genes, including multiple collagen genes as well as the EMT regulator *Runx1* by 3-15 fold, and these effects were fully attenuated by the TGFß inhibitor SB431542. Remarkably, PRDM16 expression not only repressed synthetic genes at baseline, but almost abolished the fibrotic gene response to TGFß1, resulting in a 70-100% decreased induction of *Col1a1, Col6a3, Col5a1* and *Tgfb1* (Fig. 5g). TGFß1 induced SMAD3 phosphorylation to similar levels in control and PRDM16-expressing cells, suggesting that PRDM16 represses TGFß signaling at the transcriptional, but not the receptor/signaling level (Supplemental Fig. 5j). TGFß1 treatment of control cells induced a dramatic remodeling of the actin cytoskeleton, leading to the formation of prominent stress fibers. This process was nearly completely blocked in PRDM16-expressing cells (Fig. 5h). In summary, PRDM16 is able to powerfully suppress fibrosis in an isolated, highly synthetic cellular context.

To establish how PRDM16 represses the synthetic gene program, we analyzed chromatin immunoprecipitation (ChIP) data from PRDM16-expressing fibroblasts^30^. Prominent PRDM16 binding peaks were identified at many synthetic genes that were also upregulated by *Prdm16* deletion in mouse SMCs *in vivo* and human hCaSMCs *in vitro* (Fig. 5i). These PRDM16-binding regions showed decreased levels of H3K27-acetylation upon PRDM16 expression, suggesting that they represent functional binding sites (Fig. 5i). Genes located nearby to PRDM16-binding sites that displayed PRDM16-mediated decreases in H3K27-acetylation were enriched for pathways related to fibrosis, ECM organization, and cell migration (Fig. 5j). We then performed chromatin immunoprecipitation (ChIP) for endogenous PRDM16 and H3K27Ac in mouse aortae. PRDM16 binding peaks were similarly identified at many synthetic genes such as *Ankrd1, Serpine1, Des, Tnfrsf11b, Sorbs1*, *Vim*, *Col4a1, Col8a1 and Col5a1* (Supplemental Fig. 5f-h). Motif analysis of PRDM16-binding regions identified motifs for several transcription factors, including the strongest enrichment for known PRDM16-interaction partners NR3C and EBF (Supplemental Fig. 5i). Gene ontology analysis of the PRDM16 binding regions converged on fibrotic processes, similar to the upregulated processes found in *Prdm16* KO aortae (Supplemental Fig. 5g). These results indicate that PRDM16 binds chromatin and represses transcription at synthetic gene targets.

## Discussion

This study identifies the role of the CVD-associated gene *PRDM16* as a regulator of SMC identity and atherosclerotic lesion composition. PRDM16 functions in SMCs to restrain synthetic processes and reduce synthetic SMC fate switching without affecting contractile genes or other SMC fate switches. PRDM16 is downregulated during SMC modulation in atherosclerosis, and genetic loss of PRDM16 results in the development of fibrous lesions with reduced lipid content.

The pathways that downregulate *PRDM16* expression during atherosclerosis are unknown but of substantial interest. While PRDM16 has been extensively studied in other contexts, few regulators of *PRDM16* expression have been reported. Atherosclerotic lesions are dynamic microenvironments that are influenced by many pathways. Alterations in blood flow dynamics, immune cell infiltration, ECM composition, and vascular stiffness could all contribute to *PRDM16* downregulation and the synthetic SMC phenotype transition.

The single cell analyses demonstrated that PRDM16 is required to broadly repress synthetic genes across all SMC populations, and KO mice develop highly fibrous atherosclerotic plaques. The enhanced synthetic profile and increased development of fibrous caps in *Prdm16*-deficient lesions are phenotypic features associated with a reduced likelihood of rupture in human atherosclerosis. Interestingly, we observed a stronger fibrotic phenotype in plaques from cKO (*Tagln-Cre*) compared to iKO (*Myh11-Cre-ERT2*) mice, despite a similar alteration of the gene program in both models. This may result from the longer period of *Prdm16* deficiency from development and accumulated changes in the ECM, and/or may be influenced by *Prdm16* deletion in other cell types, such as cardiac muscle, that are also targeted by *Tagln-Cre* during development^31^.

PRDM16 activates and represses gene transcription through a variety of mechanisms. In adipose and intestinal cells, PRDM16 interacts with members of the PPAR transcription factor family to promote fatty acid oxidation^16–18^. PRDM16 also promotes mitochondrial biogenesis and dynamics in adipocytes, cardiomyocytes and hematopoietic stem cells^19,32^. Interestingly, modulating *Prdm16* expression in SMCs did not affect the expression of mitochondrial or metabolic genes. These results indicate that the PRDM16-dependent repression of synthetic processes in SMCs does not involve changes in cellular metabolic programs. ChIP experiments indicate that PRDM16 binds to chromatin at putative enhancer and promoter regions of many synthetic/fibrotic genes, thereby directly repressing this gene program in SMCs. PRDM16 can repress gene transcription via interacting with various complexes, including CtBP^18^, the CBX4/polycomb complex^33^, and EHMT1/2^34–36^. Further studies will be needed to clarify the essential PRDM16-complex components and interacting partners in SMCs.

Previous studies have identified other important transcriptional regulators of SMC modulation, most notably TCF21, KLF4 and OCT4. *Tcf21* is upregulated in synthetic modulated SMCs, where it activates the expression of fibroblast and synthetic genes. *Tcf21* deletion in mouse models reduces both the number of modulated SMCs and overall lesion burden^29^. KLF4 and OCT4, both pluripotency factors, are also upregulated during SMC modulation but have contrasting roles. KLF4 suppresses contractile SMC marker expression, and promotes pro-inflammatory mediators, while OCT4 appears to maintain contractile identity and improve lesion stability^37,38^. The loss of *Klf4* or *Tcf21* results in the formation of smaller lesions, while *Oct4* deletion increases lesion size. These phenotypes are notably distinct from the effect of *Prdm16* deletion, which does not affect lesion size, but drives a selective shift of SMCs towards a more synthetic fate. Interestingly, loss of *Prdm16* upregulated the synthetic program without increasing *Tcf21* or *Klf4* expression.

SMC phenotypic switching is also associated with other vascular diseases, such as aortic aneurysms and dissections. Of note, a recent study showed that ablation of *Prdm16* aggravates elastase-induced aneurysms in mice, via increasing ADAM12 levels and inflammatory signaling^39^. We did not observe changes in *Adam12* expression in our SMC *Prdm16* KO models, nor an increase in inflammatory markers such as *Cd68* and *Lgals3*. However, PRDM16 strongly suppressed the expression of tumor necrosis factor receptor superfamily gene *Tnfrsf11b*. In atherosclerosis, TNFRSF11b regulates ECM remodeling and represses vascular calcification rather than driving inflammation^29,40^. This suggests that PRDM16 and its modulation of SMC fibrosis, may interact with other pathways, such as inflammation and matrix degradation under aneurysm-inducing conditions.

PRDM16 is also expressed in endothelial cells, though at lower levels than in SMCs. Interestingly, PRDM16 action in endothelial cells but not SMCs is required to restore arterial flow recovery following limb ischemia ^23^, suggesting that PRDM16 in SMCs is dispensable for arteriogenesis, but rather functions in mature vessels to suppress the synthetic switch. As fibrotic SMC modulation shares characteristics with endothelial-to-mesenchymal transition, we speculate that PRDM16 may also suppress this transition in endothelial cells.

In summary, PRDM16 functions as a transcriptional gatekeeper of the synthetic program in SMCs. Repressing PRDM16 function to enhance synthetic SMC activity and fibrous cap development could be beneficial for stabilizing atherosclerotic lesions and preventing lethal atherothrombotic events.

## Methods

### Mice and atherosclerosis studies

All experiments were performed according to procedures approved by the University of Pennsylvania Institutional Animal Care and Use Committee (protocol number 805649). Mice were bred and housed under the care of University of Pennsylvania University Laboratory Animal Resources. All mice for this study were male of the C57BL6/J background. The *Prdm16* floxed mouse line was generated by Patrick Seale and Bruce Spiegelman and is available from the Jackson Laboratory: *Prdm16^loxP/loxP^* (B6.129-Prdm16tm1.1Brsp/J, RRID: IMSR_JAX:024992). They were bred with *Tagln-Cre* (B6.Cg-Tg(Tagln-cre)1Her/J, Strain #:017491, RRID:IMSR_JAX:017491) or *Myh11-CreER^T^*^2^ (B6.FVB-Tg(Myh11-icre/ERT2)1Soff/J, Strain #:019079, RRID:IMSR_JAX:019079). Floxed animals without Cre were used as controls. Animals were raised at room temperature on standard chow (LabDiet, 5010) with a 12-h light– dark cycle at room temperature (22 °C) unless specified otherwise. For atherosclerosis experiments, mice were placed at thermoneutrality (30 °C) at weaning (21 days old) and kept at thermoneutrality until the end of the experiment. For iKO studies, tamoxifen (Sigma, T5648; stock 20 mg/mL in corn oil) was injected intraperitoneally at a dose of 100 mg/kg for 4 consecutive days. For atherosclerosis studies, the mice were retro-orbitally injected at 8 weeks with AAV8-PCSK9 D377Y (Vector Biolabs) at a dose of 5 x 10^11^ GC/mouse, as described by to Bjørklund et al.^41^. The following day, the mice were put on Western Diet composed of 40% fat and 0.15% cholesterol (Research Diets, D12079B) for 12 weeks, or where indicated, 18 weeks. The mice were weighed bi-weekly, and plasma samples were acquired every 4 weeks for lipid measurements after a 3-hour fasting period.

### Histology, Microscopy and Image Analysis

At 20 weeks, the mice were sacrificed, and the aorta was perfused through the left ventricle with phosphate-buffered saline. The aortic arch was isolated and fixed in 1% paraformaldehyde for 16 hours, followed by 4 PBS washes and sequential dehydration and paraffin-embedding. Longitudinal sections (6 µm), and a series of 4-5 subsequent sections through the aorta of 24 µm apart were stained with hematoxylin and eosin, imaged on a Keyence inverted microscope and analyzed for average plaque area per aorta. A Verhoeff van Gieson staining kit (Polysciences) was used to visualize elastin fibers, and Sirius-red staining, counterstained with Fast-Green, to analyze collagen content. Slides were imaged on a Zeiss Axio Observer 7 inverted microscope, and collagen positive area per plaque area was selected with the “color range” option in Adobe Photoshop and calculated as pixels per lesion area. For immunohistochemistry, slides were deparaffinized and heat-induced antigen retrieval was performed in Bull’s Eye Decloaking buffer (Biocare). Primary antibody incubation was done overnight, stained with secondary antibody, and then developed with tyramide signal amplification (Akoya Biosciences). For a list of antibodies and concentrations, see supplemental table 1. Immunohistochemistry stainings were imaged on a Zeiss Axio Observer 7 or a Leica SP8 Confocal Microscope.

### Oral glucose tolerance test (OGTT) and Blood Pressure

Mice were fasted for 16 hours, before receiving glucose (2 g/kg) by oral gavage. Blood glucose levels were measured directly from tail vein blood at 0, 15, 30, 45, 60 and 90 min using an Ascensia Contour Next glucometer (Medline, USA). For blood pressure measurements A Visitech BP-2000 series II arterial pressure analysis system was used. At 8 weeks old, the mice were trained for 3-5 subsequent days at the same time until consistent measurements were obtained. Each day, 3-10 individual measurements were recorded and averaged, for a total of 5 days. Measurements were excluded if the mouse could not get consistent reads for at least 3 measurements.

### Cell culture, migration assays and analysis

hCaSMCs were immortalized as previously described ^41^. Transcriptional activation of PRDM16 was achieved using the synergistic activation mediator (SAM) complex. guide RNAs (gRNA)s were designed using CRISPick ^42^, CHOPCHOP ^43^, and the guidelines outlined by Nageshwaran et al. ^44^. The gRNA (5’-GCAATCTGACACCCCTCGCCG-3’) was cloned into a SAM vector using golden gate cloning. Proper insertion of the gRNA into the vector was verified using Sanger sequencing. hCASMC-hTert were then transduced concomitantly with the lentiSAMv2 and the lentiMPHv2 (both vectors were a gift from Feng Zhang (Addgene plasmid # 75112 and # 89308) in 8 ug/ml polybrene (Millipore, TR1003-G). Transduced cells (hCASMC-PRDM16 CRISPRa) were selected using Blasticidin and Hygromycin, respectively at 10ug/ml and 400 μg/ml (ant-bl-05, InvivoGen; 10687010, Gibco) and maintained in SMC basal medium supplemented with a complementPack (C-22162, PromoCell). The expression of PRDM16 was measured via qPCR and compared to HCASMC-hTert transduced with both lentiSAMv2 and lentiMPHv2, the former vector being empty (HCASMC-EV). For lentiviral overexpression, a HA-tag was added to the *Prdm16* cDNA (CCDS71532) and cloned into pLentiV2-CMV Puro using New England Biolabs HiFi, replacing the GFP CDS. pLenti CMV GFP Puro (658-5) was a gift from Eric Campeau & Paul Kaufman (Addgene plasmid # 17448). The empty vector was used as a control (Addgene plasmid # 17448). hCaSMCs were infected with 10x concentrated lentivirus in the presence of 8 µg/mL polybrene overnight, and selected with 2.5 µg/mL puromycin. BMC fibroblasts^35^ were infected with the indicated retroviruses in the presence of 8 µg/mL polybrene, and positive clones were selected with 3 µg/mL puromycin. Where indicated, Cells were treated with recombinant mouse TGFß-1 (R&D Systems) at a final concentration of 5 ng/mL, and SB431542 (Invivogen) at a final concentration of 10 µM. DMSO was used as a vehicle control. For migration assay, cells were seeded to confluency on µ-slides (Ibidi), coated with 5 µg/mL fibronectin (Corning). The next day, the cells were pre-treated for 2 hours with 2 µM Mitomycin C (Sigma). A scratch was created, the cells were washed, and scratch closure was observed in an EVOS microscope every 10 minutes for 14 hours at 37°C and 5% CO_2_. Scratch area was quantified every 2 hours using ImageJ software. For Proliferation measurements, cells were seeded at 10% confluency, grown for 72 hours and cell density was assessed using 3-(4,5-Dimethylthiazol-2-yl)-2,5-Diphenyltetrazolium Bromide MTT (ThermoFisher).

### RNA isolation, RT-qPCR and bulk RNAseq analyses

Total RNA was extracted using TRIzol (Invitrogen). For *in vitro* experiments, the PureLink RNA columns (ThermoFisher) were used, and for mouse tissuses a TRIZol/chloroform extraction was performed according to the TRIzol manual. RNA concentrations were measured with the Nanodrop (ThermoFisher), and cDNA was synthesized using the MultiScribe Reverse Transcriptase kit (Thermo Scientific). Real-time quantitative PCR was performed on a ABI7900HT PCR machine with SYBR green (Applied Biosystems). Fold changes were calculated using the ΔΔCT method using TATA-box binding protein (Tbp) as housekeeping gene. For Bulk RNAseq analyses, the aorta was isolated, freed of PVAT and adventitia, cut open and the endothelial layer was gently brushed off. RNA was then isolated using the PicoPure RNA isolation kit (ThermoFisher). Next-generation sequencing (NGS) using Illumina NGS technology was generated by Genewiz using the NovaSeq 6000 analyzer. Fastq files were aligned to GRCm39 using kallisto’s (version 0.44.0) quant function with default settings. The abundance.tsv files were read into R (version 4.3.0) with the package tximport (version 1.28.0) and the parameter “txOut = F”, to get gene level quantification. The ComBat_seq function from the package sva (version 3.35.2) was utilized to correct the raw counts expression matrix from batch effects observed from separate sequencing runs. The corrected expression matrix was converted to a DGE object using the package edgeR (version 3.42.4) in which lowly expressed genes were filtered out, normalization factors were calculated to account for differences in sequencing depth across samples using a trimmed mean of M-values (TMM), and this data was converted to log 2 counts per million (CPM). This batch corrected adjusted matrix would be used for heatmap visualizations. For differential gene expression, the raw un-batch corrected data was first filtered out for lowly expressed genes across samples and the package limma (version 3.56.2) was used for identifying differentially expressed genes across conditions. The model matrix was designed to take both condition and sequencing run into account for the design. The data was converted using voomWithQualityWeights to apply the voom transformation and to account for variations in individual samples. Then a linear model was fit to the data with a robust M-estimation to minimize the effect of outliers. Then specific group to group comparisons were made based on conditions, these contrasts were applied to the fitted model, and then empirical Bayes moderation was applied to the data to calculate statistics. Minus-average plot (MA) was constructed utilizing ggplot2 (3.5.1). Gene set enrichment analysis was performed using the fgsea package (version 1.26.0) on the differentially expressed gene list which was ordered by “-log10(p-value)*sign(logFC)” with the reference gmt files obtained from MSigDB (https://www.gsea-msigdb.org/gsea/msigdb/human/collections.jsp). Functional enrichment analysis was performed using the gprofiler2 package (version 0.2.3) in which a subset of genes were used for the analysis. Heatmaps were created using the ComplexHeatmap package (version 2.16.0) where the data was converted to Z-score for scaling.

### Single Cell RNAseq

scRNAseq was performed using the 10x Flex Platform using the Chromium Next GEM Single Cell Fixed RNA Sample Preparation Kit (1000414). The aortae were dissected from root until ileac bifurcation, washed twice in cold RPMI, excess liquid was removed and the tissue was sliced in 1 mm rings on a chilled glass slide and fixed in 1 mL fixation buffer / 25 mg tissue (4% PFA along with 10x conc fix perm buffer) for 16h at 4 °C without agitation. The next day, the tissue pieces were centrifuged for 5 min @ 850 x g, and washed in 2 mL cold PBS. The tissue was then resuspended in 1 mL quenching buffer with 0.1 volume of Enhancer, and a final glycerol concentration of 10% and stored at −80 °C. The aortae were digested separately in 2 mL pre-warmed RPMI 1640 with 1 mg/mL of Liberase TM (Roche 05401020001) on a Miltenyi GentleMACS at 37 °C using the following program: (spin 50 rpm for 20 min, [spin 1000 rpm for 30s, spin −1000 rpm for 30s (2x)].The digested tissues were pooled and strained using a 30 µm strainer, and the tube and filter were rinsed with PBS. The cells were centrifuged at 1200 x g for 10 min, and the sample was resuspended in 250 µL quenching buffer. 5% was taken out for DAPI staining and counting. Probe hybridization and cell capture was performed using the 10x Chromium Fixed RNA Kit, Mouse Transcriptome kit (1000495) according to the manufacturer’s instructions except for an increase in centrifugation speed (1200 x g for 10 min). Custom probes were designed using the instructions by 10x and probes were obtained as oPools by IDT DNA Technologies.

### Single cell analysis

#### cKO scRNAseq processing

Fastq files were aligned to the mouse genome (mm10) with cellranger (version 7.2.0) using the command cellranger multi which was aligned to a customized chromium mouse transcriptome probe set (version 1.0.1) to include manually designed probes for Prdm16 Exon 9 detection. The raw matrix files for the Tagln Control and Tagln Prdm16 Exon 9 Knockout were read into R (version 4.3.0) and processed with DropletUtils (version 1.20.0) where droplets with a false discovery rate less than or equal to 0.01 were retained. These new filtered datasets were then processed with SoupX (version 1.6.2) using it’s autoEstCont and adjustCount features to estimate the contamination fraction and then adjust the expression matrices. Taking the newly created ambient RNA expression corrected matrices, we used seurat (version 4.3.0) for quality control processing, filtering, and integration. The function CreateSeuratObject with the parameters “min.features = 20, min.cells = 100” was used to make seurat objects. Then, light filtering of the datasets were performed with the following parameters “nCount_RNA > 500 & nCount_RNA < 20000 & nFeature_RNA > 150 & nFeature_RNA < 8000 & percent.mt < 10”. Then each samples counts were scaled for each cell by the total number of molecules detected and then multiplied by 10000 and log transformed. The top 2000 most variable features were found in each dataset with a “vst” selection method and the data had its expression scaled and centered for each gene across all cells while regressing on the mitochondrial percentage. Principal component analysis (PCA) was performed where dimensions 1 through 15 were utlized for both datasets as encompassing a majority of the variation in the data. Using these dimensions, a k-nearest neighbor graph was constructed and clustering was performed on the nearest-neighbor graph using a Louvain algorithm and a resolution of 0.2. A uniform manifold approximation and projection (UMAP) was created for visualization purposes. The two processed seurat objects were run through scDblFinder (version 2.0.3) where cells were identified as singlets or doublets.

After identifying all cells that were possible doublets, we re-created the starting seurat objects with the same parameters, retained all cells that were identified as singlets, and then re-filtered the data with stricter quality control parameters of “nCount_RNA > 500 & nCount_RNA < 10000 & nFeature_RNA > 500 & percent.mt <= 1” for both datasets. Then each samples counts were scaled for each cell by the total number of molecules detected and then multiplied by 10000 and log transformed. The top 2000 most variable features were found in each dataset with a “vst” selection method. The top 2000 variable genes shared across datasets were selected and common reference points were identified between datasets using a canonical correlation analysis (CCA) and log normalization method. The data was then integrated together and the subsequent dataset’s expression was scaled and centered for each gene across all cells while regressing on the mitochondrial percentage, S.Score, and G2M.Score calculated from cell cycle scoring. Harmony (version 1.2.0) was run on the integrated data for batch-effect correction but it’s effects appeared neglible so the PCA was utilized for subsequent processing. Principal components 1 through 10 were utlized, a k-nearest neighbor graph was constructed and clustering was performed on the nearest-neighbor graph using a Louvain algorithm and a resolution of 1.0. A UMAP was created for visualization purposes and UMAP projections were constructed using seurat’s DimPlot function, gene expression patterns were plotted using FeaturePlot function, and changes in percentage and expression levels in cells using DotPlot. For cluster classification, the top expressed genes were identified for each cluster using seurat’s FindMarkers function using a Wilcoxon Rank Sum test and these genes were cross-referenced against the literature and online databases to annotate clusters.

After identifying all cells that were possible doublets, we re-created the starting seurat objects with the same parameters, retained all cells that were identified as singlets, and then re-filtered the data with stricter quality control parameters of “nCount_RNA > 500 & nCount_RNA < 10000 & nFeature_RNA > 300 & percent.mt <= 1” for both datasets. Then each sample’s counts were scaled for each cell by the total number of molecules detected and then multiplied by 10000 and log transformed. The top 2000 most variable features were found in each dataset with a “vst” selection method. The top 2000 variable genes shared across datasets were selected and common reference points were identified between datasets using a canonical correlation analysis (CCA) and log normalization method. The data was then integrated together and the subsequent dataset’s expression was scaled and centered for each gene across all cells while regressing on the mitochondrial percentage. Harmony (version 1.2.0) was run on the integrated data for batch-effect correction but it’s effects appeared neglible so the PCA was utilized for subsequent processing. Principal components 1 through 10 were utlized, a k-nearest neighbor graph was constructed and clustering was performed on the nearest-neighbor graph using a Louvain algorithm and a resolution of 1.0. A UMAP was created for visualization purposes and UMAP projections were constructed using seurat’s DimPlot function, gene expression patterns were plotted using FeaturePlot function, and changes in percentage and expression levels in cells using DotPlot. For cluster classification, the top expressed genes were identified for each cluster using seurat’s FindMarkers function using a Wilcoxon Rank Sum test and these genes were cross-referenced against the literature and online databases to annotate clusters.

#### ChIP-sequencing

PRDM16 and H3K27ac ChIPseq was performed by Kissig et al. ^30^ on fibroblasts expressing a control construct (puromycin resistance) or Prdm16. Target enrichment was calculated as percent input. ChIP-seq reads for Prdm16 and H3K27Ac were aligned to mouse genome, mm9, and further processed for peak-calling and genome browser track creation as described^30,45^. Data is available through GSE86017. For PRDM16 ChIP in C57Bl6/J aortae, 12 were dissected, dual-crosslinked with 1.5mM EGS for 20 min followed by 1% formaldehyde for 10 min, then washed and snap frozen in liquid nitrogen. Aortae were pooled into a 2mL safe-lock tube with a 3mm stainless steel bead and cryo-milled into a powder using 2 cycles of 5Hz for 30 sec and 23 Hz for 30 sec. The pulverized aortas were placed on wet ice, allowed to thaw slightly and then resuspended in 0.6% SDS lysis buffer. Chromatin was sheared using a microtip probe sonicator at 25% Amplitude, 30 sec on, 20 sec off for 5 minutes. After the first minute of shearing the insoluble ECM clump was removed with a pipet tip and shearing resumed for the remaining 4 minutes. Dilution buffer was added to make a 0.1% SDS final IP concentration and the sample was split into 3 separate immunoprecipitations with approximately 10ug chromatin per IP. Antibodies are listed in extended material table 2. Samples were incubated with the indicated antibody concentrations at 4 °C overnight and Protein A/G magnetic beads were used to capture target chromatin for subsequent washing and elution. DNA was reverse crosslinked overnight at 65 °C with RNase A and treated with proteinase K for 2hrs at 50 °C. ChIP DNA was purified using Active Motif Chromatin IP Purification Kit and used for downstream library prep. Sequencing libraries were made according to manufacturer’s recommendations using the Roche KAPA HyperPrep Kit and Roche Universal Dual-Index Adapters. Size selection was performed using Beckman Coulter SPRI Select beads for an 800 bp upper limit cut-off.

#### ChIPseq Analysis

Fastq files were first trimmed using the package trim_galore (version 0.6.10) with default settings and the “--paired” parameter selected. Paired trimmed reads were aligned to the mouse genome (mm10) using bowtie2 with the following parameters “-N 1” and this SAM output was piped into samtools view (version 1.6) with the parameters “-bSF4”. This BAM file was then sorted, duplicate reads were removed, reads that could not be uniquely mapped were dropped, and the file was indexed. The processed BAM file for the specified sample was then converted to bigwig files using bamCompare from deeptools (version 3.2.2) comparing to the IgG sample with the specifications of “--binSize 20 --smoothLength 60 –extendReads 150 --effectiveGenomeSize 2653783500 --operation subtract” for the H3K27ac sample. The same parameters were run for the Prdm16 sample with added specifications of “--centerReads” selected. HOMER (version 4.11) was used for peak calling where tag directories were first created for each sample utilizing makeTagDirectory with the parameters “-single -genome mm10” specified. The function findpeaks was used for peakcalling with parameters “-size 200 -F 2 -L 2 -fdr 0.01 -center -region” specified for Prdm16 compared to the IgG input tag directory. The same parameters were run for IgG without calling “-i -center” and “-region” and the H3K27ac sample was run with “-style histone”. To find known and de novo motifs, Homer’s findMotifsGenome.pl function was run with parameters “-dumpFasta -mask”. Homer’s annotatePeaks.pl was used to annotate the called peaks. The fasta output from Homer, along with the motifs, were utilized in MEME Suite (version 5.5.7) in programs tomtom and DREME which were run with default settings. IGViewer was used to visualize tracks.

#### Immunoblotting

Total cell lysates were prepared in RIPA buffer (150 mM NaCl, 1% NP-40, 0.1% sodium deoxycholate, 0.1% SDS, 100 mM Tris-HCl, pH 7.4) supplemented with protease inhibitors (Roche), and phenylmethylsulfonyl fluoride (Sigma). Lysates were sonicated and 4x NuPAGE LDS Sample Buffer (Thermo Scientific) with 10% β-mercaptoethanol was added. After denaturation at 95 °C for 5 minutes, the denatured proteins were separated on NuPAGE Novex 4–12% Bis-Tris gels (Thermo Scientific) and transferred to Amersham Protran nitrocellulose membranes (Sigma) and probed using horseradish peroxidase-conjugated secondary antibodies (Cell Signaling Technologies) and SuperSignal West Pico PLUS Chemiluminescent Substrate (Thermo Scientific). See supplemental table 1 for a list of primary antibodies.

**Supplemental Table 1:** Antibodies used for Immunohistochemistry and Immunoblotting.

| Antibodies used for immunohistochemistry |  |  |  |
| --- | --- | --- | --- |
| ACTA2 | 1:1000 | Sigma | A5228 |
| TAGLN | 1:500 | Abcam | Ab10135 |
| DAPI | 1:1000 | Sigma | D5942 |
| Ki67 | 1:250 | Abcam | ab16667 |
| Lumican | 1:200 | Novus | NBP2-76847 |
| MYH11 | 1:400 | Abcam | ab224804 |
| CD68 | 1:200 | Abcam | ab955 |
| PRDM16 | 1:500 | Made in house |  |
| Perilipin 2 | 1:200 | Novus | NB110-40877 |
| PECAM-1 | 1:200 | HistoBioTec | DIA-310 |
| Antibodies used for ChIP |  |  |  |
| PRDM16 | 6 μg | Made in house | |
| H2K27ac | 2 μg | Active Motif | 39133 |
| Antibodies used for Immunoblotting |  |  |  |
| PRDM16 | 1:500 | Made in house |  |
| pSmad3 | 1:1000 | Cell Signaling Technologies | #9520 |
| Smad3 | 1:1000 | Cell Signaling Technologies | #9523 |
| Actin | 1:1000 | Milipore | MAB1501 |

#### Statistical Analysis

comparisons between two groups, a 2-sample t-test was employed, while comparisons among multiple groups with single or two factors were assessed using 1 or 2-way ANOVA, respectively when passing a Kolmogorov-Smirnov test for normality. When the requirements for normal distribution were not met, samples were analyzed using a Kruskall-Wallis test. To account for multiple comparisons, the Holm-Šídák correction was applied where appropriate. Prior to analysis, the ROUT method was used to identify and remove outliers. All data points represent individual, not repeated measurements. Wound closure rates were fitted using a simple linear regression analysis and an F-test for Regression Analysis was used to assess significance. All statistical tests were performed in GraphPad Prism V10.

#### Analysis of human bulk RNAseq and scRNAseq

For bulk RNAseq of early- and advanced lesions, human carotid plaques of 38 patients (10 female, 28 male) were prepared, sequenced, and the paired analysis was performed as described by Fidler, et al. (2024)^25^. The scRNAseq analysis on 21 coronary endarterectomy samples was performed as described by Bashore et al. (2023)^46^. Analyis, QC and visualization of the feature plots was performed using the Scanpy package in Python.

#### Re-analysis of public scRNAseq and snATACseq datasets Single Cell RNA Sequencing

All figures generated for the single cell RNA sequencing dataset were generated using data generated by Wirka et al.^29^, GEO accession GSE131776. The processing pipeline used followed the methods described in the paper. The raw fastq files were obtained using the sratoolkit (version 3.0.6) using the prefetch and fasterq-dump commands. The single cell RNA sequencing data was preprocessed using the 10x Genomics pipeline with Cell Ranger (version 7.2.0), using the reference mouse genome with default parameters. Samples from each patient were analyzed separately and the cellranger output files were read into R for further downstream processing.

The Seurat package (version 4.3.0) was used for quality control and clustering. Samples that expressed fewer than 500 genes and genes expressed in fewer than 5 cells were removed from the dataset. The samples were then merged utilizing the merge function. Cells with a mitochondrial percentage greater than 7.5 were removed as indication for dead or dying cells. Cells with a more than 3500 genes detected were discarded as they were indicative of “doublets”. The remaining cells underwent library size normalization using a log normalization method and a scaling factor of 10000. The top 2000 variable features were selected using a variance stabilizing transformation (vst) method and the data was scaled and centered. Principal component analysis (PCA) for dimensionality reduction was performed, followed by clustering in PCA space using a graph-based clustering approach. Dimensions 1 through 30 were retained in the dimensionality reduction and a resolution of 0.3 was used when finding clusters. The t-distributed stochastic neighbor embedding was then used for two-dimensional visualization of the resulting clusters. Plots were generated using Seurat’s DimPlot for the two-dimensional visualization and FeaturePlot for gene expression on the two-dimensional layout. The cluster-specific genes (marker genes) for each cluster were identified with the Seurat default method, a Wilcoxon rank sum test.

#### Single Nucleus ATAC Sequencing

All figures generated for the single nuclei ATAC sequencing dataset were generated using data generated by Turner et al.^15^, GEO accession GSE175621. The raw fastq files were obtained using the sratoolkit using the prefetch and fasterq-dump commands. The single nuclei ATAC sequencing was preprocessed using the 10x Genomics pipeline with Cell Ranger ATAC (version 7.2.0), using the reference GRCh38 genome with default parameters. Samples from each patient were analyzed separately and the cellranger output files were read into R for further downstream processing.

The cell ranger files were processed in R utilizing Seurat and Signac (version 1.10.0). A unified peak set for all samples was created utilizing the granges function from the GenomicRanges package (version 1.52.0). This unified peak underwent an inter-range transformation with the reduce function from Signac, was filtered for non-standard chromosomes with a coarse pruning mode, blacklisted regions in hg38 were removed, and peaks with a width smaller than 20 and larger than 10000 were removed. After creating the unified peak set among all samples, a matrix of peaks per cell, the chromatin assay object and seurat object for each sample were made using the commands FeatureMatrix, CreateChromatinAssay, and CreateSeuratObject. The snATAC-seq reads from different individuals were combined at the single-nucleus level and nuclei were filtered out that had a TSS enrichment < 0.5 and > 6, total number of fragment counts per cell > 60000, percent reads in peak < 1, nucleosome signal > 2, and a total number of fragments in peaks > 30. We then computed the term-frequency inverse-document-frequency on the combined dataset with default parameters and found the top features with a lower percentile bound of ‘q0’. A singular value decomposition was applied to the data. The dimensionality reduction was conducted using a latent semantic indexing algorithm with omitting the first principal component as this component captures sequencing depth rather than biological variation, therefore dimensions 2 through 15 were used for cell clustering. Cell clusters were identified by a shared nearest neighbor (SNN) modularity optimization-based clustering algorithm from Seurat. The algorithm used was the smart local moving algorithm (SLM) for its enhanced efficiency for community detection in large networks with a resolution of 0.1.

#### Integrating Single Nuclei ATAC Sequencing data with Single Cell RNA Sequencing data

Integration of the single nuclei ATAC sequencing dataset (Turner et al.) with the single cell RNA sequencing dataset (Wirka et al.) to obtain clustering by vascular cell type was performed using the FindTransferAnchors and TransferData functions in the Seurat package. Prior to the anchor transferring steps, genes in the single cell RNA sequencing mouse dataset were converted to their orthologous versions in common with humans. This was done using the package biomaRt (version 2.56.1) and utilizing the GRCm39 mouse genes and their human ortholog equivalents. After conversion of mouse to human genes, the single cell RNA sequencing dataset was used a reference to overlay the single nuclei ATAC sequencing dataset onto. A canonical correlation analysis (CCA) was used for the reduction in finding transfer anchors, dimensions 1 through 50, and the top 2000 variable features in the reference dataset were used. After, the data was transferred between the two datasets using dimensions 2 through 50, a latent semantix indexing (lsi) reduction, a k.weight of 10, and using the annotated cell markers established in the single cell dataset to transfer onto the single nuclei data. The peak expression plots were generated from the Signac package using the CoveragePlot function with window size of 250. The genome track was generated using the package Gviz (version 1.44.1) using the hg38 genome. The vlnplots to show gene expression levels for each cell type were created using the dittoPlot function from the dittoSeq package (version 1.12.1).

## Data Availability

Bulk RNA sequencing data will be deposited in the GEO before publication, after which the accession number will be provided. The human coronary artery snATACseq data from Turner et al.^10^ is available through GSE175621. The Human coronary artery scRNAseq is from Bashore et al. (Fig 1, available through GSE253904^11^. Human scRNAseq Human coronary artery scRNAseq (Extended Fig. 1 is described in Paloschi et al. and ^12^ is available through GSE247238. bulk RNAseq of early and late-stage lesions by Fidler et al.^7^ is available through GSE248395. ChIPseq data from Kissig et al.^5^ is available through GSE86017.

## Acknowledgements

We thank the members of the Seale lab, the cell and developmental biology department (CDB) and the institute for diabetes, obesity and metabolism (IDOM). We are grateful to Soon Tang, Ioana Soaita, Matthew Gavin (UPenn), Mei Zhang and the single cell technology core (Childrens Hospital of Philadelphia), and Cindy van Roomen (Amsterdam UMC) for help with experimental design. We also thank Jeff Ishibashi for editing the manuscript. This work was supported by NIH grants DK123356, AR078650, and DK120982 to P.S.; R01HL164577 and R01HL148239 to CLM. MPR and CX were supported for this work by NIH awards R01HL150359, R01HL166916, and R01HL169766. J.M.E.T. was supported by a Rubicon grant 40-45200-98-21118 from the Netherlands Organization for Health Research and Development).

## Author Contributions

JMET and PS jointly conceptualized the project and wrote the manuscript. JMET performed the experiments, imaging, data analysis and graphical editing. LC processed tissue sections for histology and performed immunostaining. RPC performed bioinformatic analyses. AHW performed ChIP experiments. SF, KD, and EB assisted with experiments. HWL analyzed ChIP data. CLM and GA provided human coronary artery smooth muscle cells engineered to stably express PRDM16. CX and MR provided the human scRNAseq data. HW and LM provided the human plaque bulk RNAseq and human scRNAseq. EL assisted in study design and edited the manuscript.

## Materials and Correspondence

Further information and requests for resources and reagents should be directed to Patrick Seale.

