## Supplemental Figures for "PRDM16 controls smooth muscle cell fate in atherosclerosis"

### Supplemental Figure 1

**a**

*Coronary Artery Disease*

| Variant | Reported.Trait | p.value | OR | Beta | CI | Study.Accession |
|---|---|---|---|---|---|---|
| rs2493298 | Coronary artery disease | 2 x 10-9 | NA | 0.0567 unit incr. | 0.038-0.075 | GCST005196 |
| | Coronary artery disease | 2 x 10-11 | 1.05 | NA | 1.035-1.065 | GCST90132314 |
| | RA-system medication use | 2 x 10-11 | NA | 0.058988318 unit incr. | 0.042-0.076 | GCST007930 |
| | RA-system medication use | 1 x 10-11 | NA | 0.059 unit incr. | 0.042-0.076 | GCST90018988 |
| | Coronary artery disease | 2 x 10-9 | NA | 0.057 unit incr. | 0.037-0.077 | GCST005194 |
| | Coronary artery disease | 1 x 10-9 | NA | 0.0514 unit incr. | 0.035-0.068 | GCST005195 |
| | Coronary artery disease | 2 x 10-9 | NA | 0.0567 unit incr. | 0.038-0.075 | GCST005196 |
| | Coronary artery disease | 2 x 10-11 | 1.05 | NA | 1.035-1.065 | GCST90132314 |
| | RA-system medication use | 2 x 10-11 | NA | 0.058988318 unit incr. | 0.042-0.076 | GCST007930 |
| | RA-system medication use | 1 x 10-11 | NA | 0.059 unit incr. | 0.042-0.076 | GCST90018988 |
| | Coronary artery disease | 2 x 10-9 | NA | 0.057 unit incr. | 0.037-0.077 | GCST005194 |
| | Coronary artery disease | 1 x 10-9 | NA | 0.0514 unit incr. | 0.035-0.068 | GCST005195 |
| rs7413494 | Coronary artery disease | 1 x 10-9 | 0.965 | NA | 0.955-0.976 | GCST90132314 |
| | Myocardial infarction | 5 x 10-8 | NA | 0.0553 unit incr. | 0.036-0.075 | GCST90018877 |
| | Coronary artery disease | 1 x 10-9 | 0.965 | NA | 0.955-0.976 | GCST90132314 |
| rs2493296 | Systolic blood pressure | 2 x 10-20 | NA | NA | NA | GCST007087 |
| | Cardiovascular disease | 3 x 10-8 | NA | NA | NA | GCST007072 |
| rs12735779 | Cardiovascular disease in type 2 diabetes | 8 x 10-6 | 3.72 | | 2.09-6.63 | GCST011371 |
| rs2455132 | Small vessel stroke | 1 x 10-8 | 1.1 | NA | 1.06-1.13 | GCST90104537 |

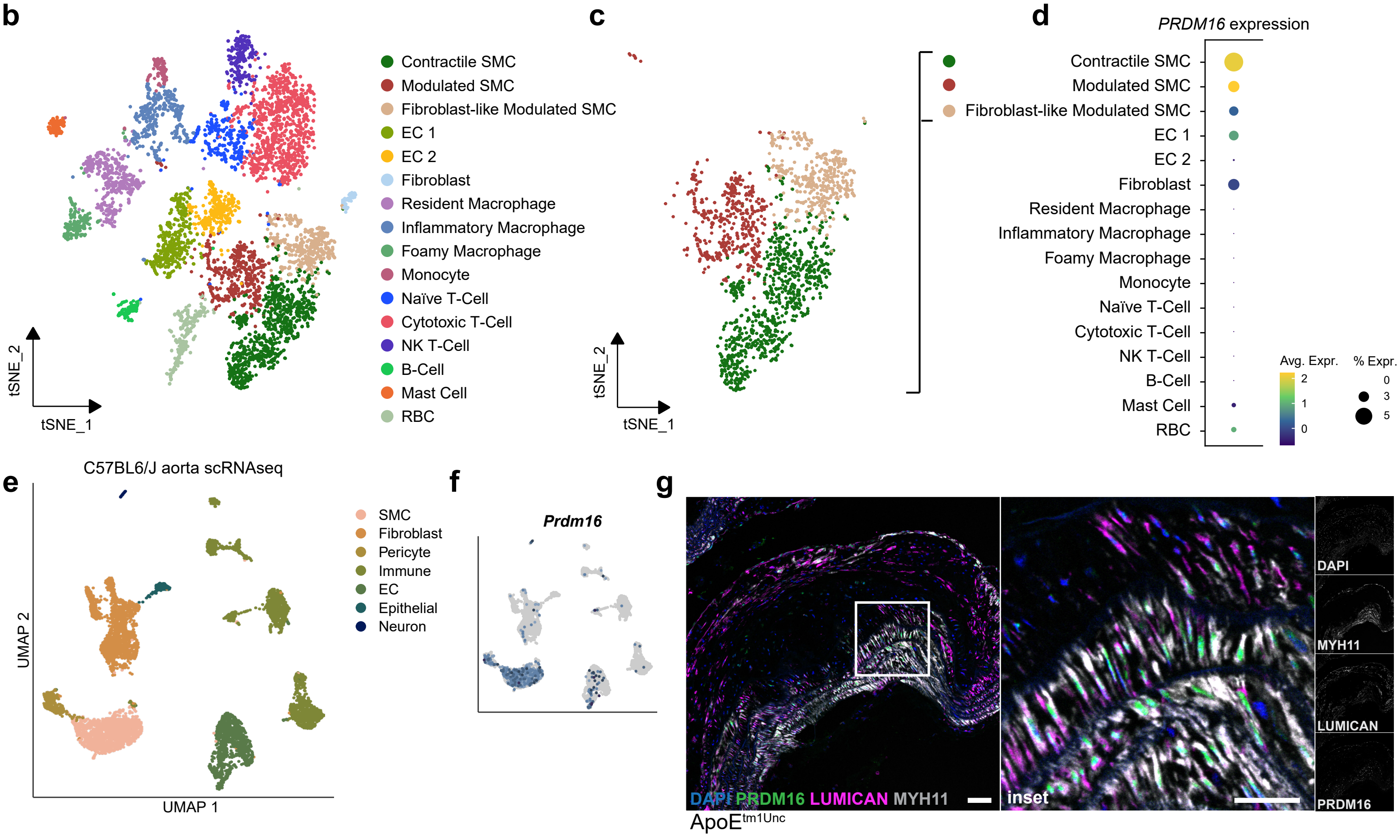


**Supplemental Figure 1 | a,** Variants in the *PRDM16* gene associated with CAD and CVD, retrieved from the NHGRI-EBI GWAS catalog. **b,** tSNE showing the major cell clusters identified from scRNAseq of human carotid artery plaques n=9. **c,** Subset of the three SMC clusters. **d,** *PRDM16* expression in all clusters from (b). **e,** UMAP of gene expression in 6,753 cells from adult thoracic aortae of 13-week-old male CD1 mice[28] **f,** UMAP feature plot of *Prdm16* expression in clusters from (e). g, Immunostaining for PRDM16 (green), MYH11 (white), Lumican (magenta) and DAPI (nuclei, blue). In a representative sagittal section of an aortic lesion with a fibrous cap from *Apoe-/-* mice fed a western diet for 18 weeks. Scale bar: 50 μm.

### Supplemental Figure 2

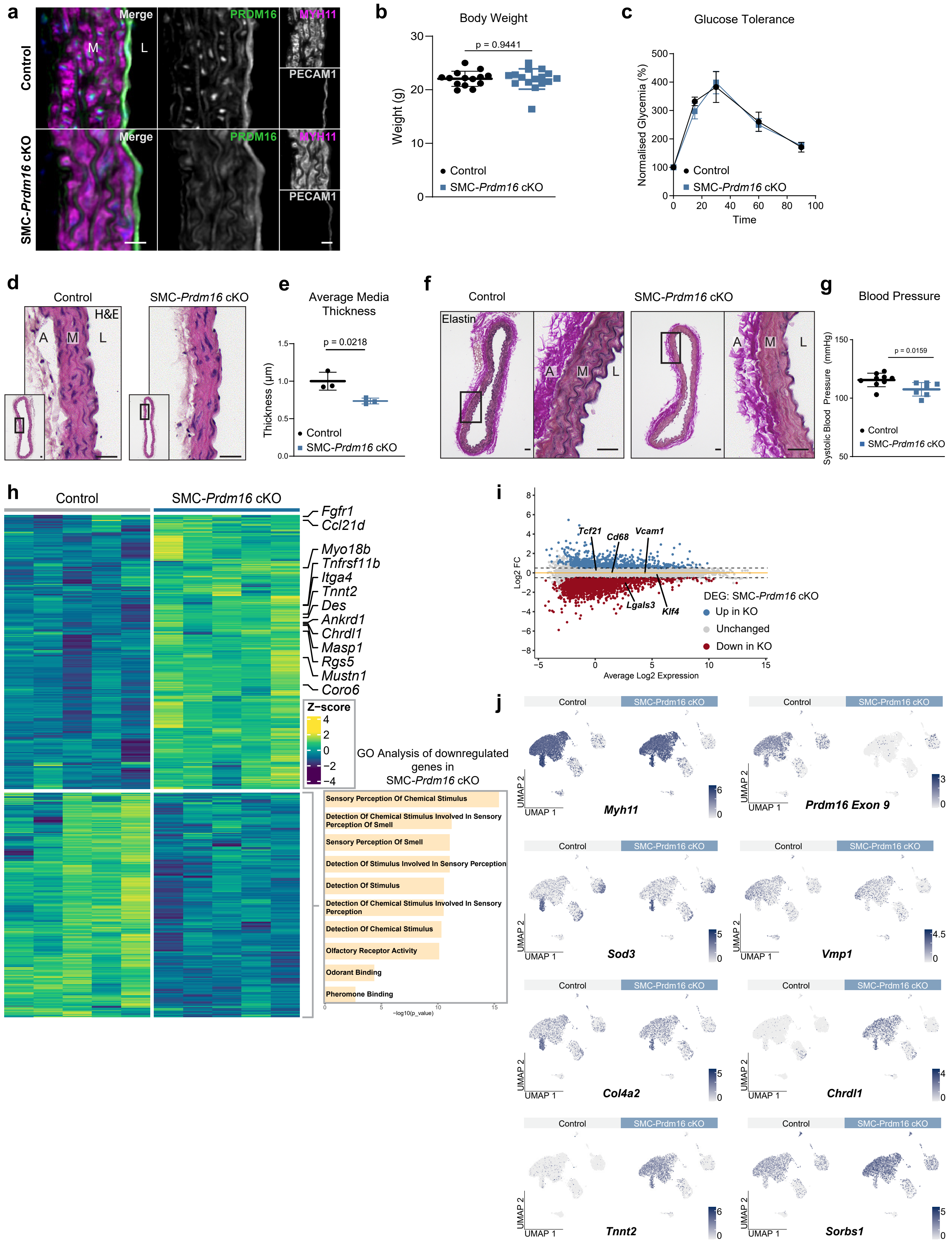

#### Supplemental Figure 2

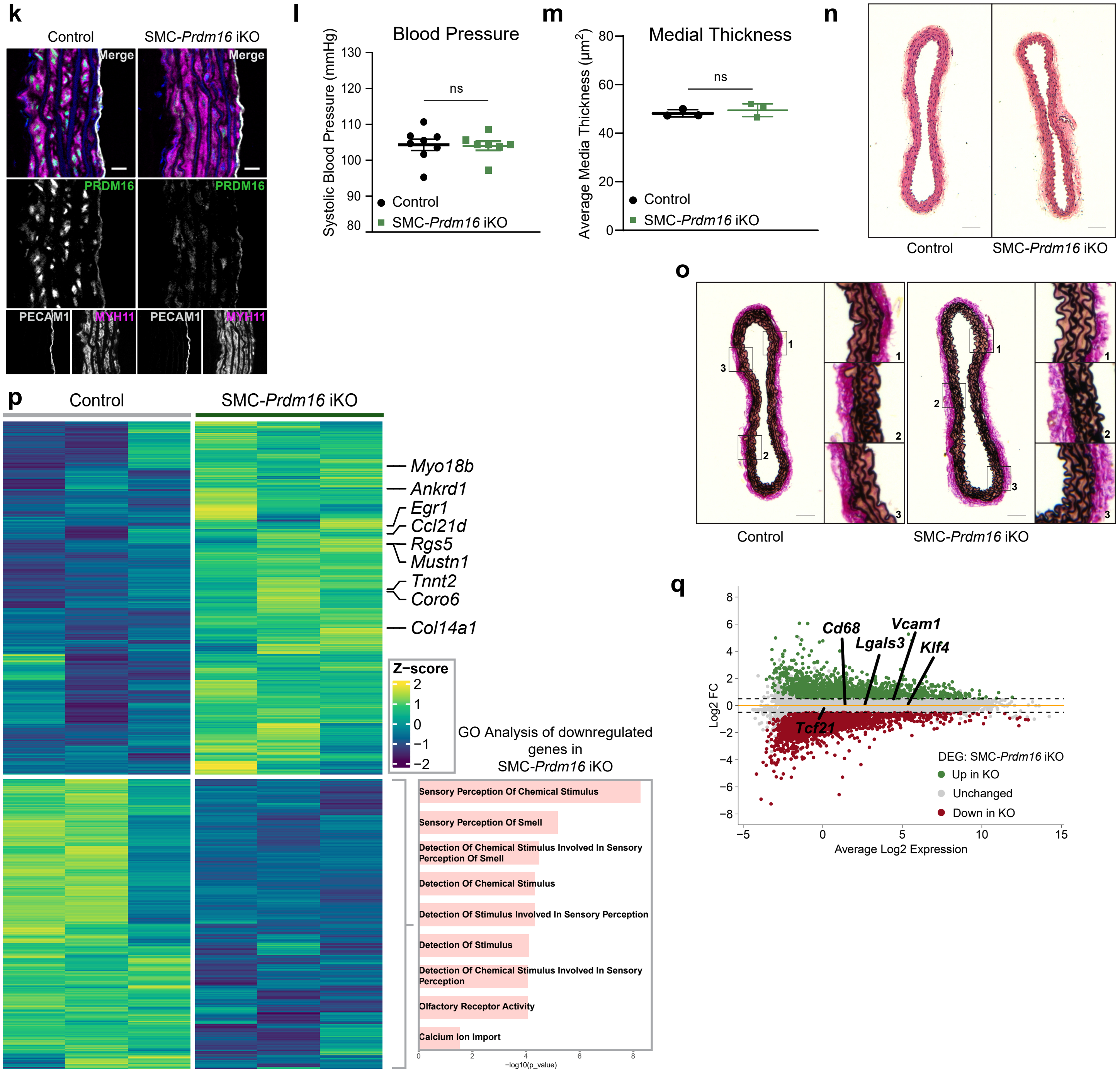


**Supplemental Figure 2 | a,** Immunostaining for MYH11 (magenta), PRDM16 (green), PECAM-1 (CD31, white), and DAPI (nuclei, blue), and in cross sections of descending aorta from control and and cKO mice. Representative of n=3/3 Ctrl/KO. Scale bar: 20 μm. M: tunica media, L: lumen. **b,** Bodyweight of 8-week-old male control and cKO mice. Values are shown as mean ±SD, n=14 control vs 16 cKO. **c,** Oral glucose tolerance test in 8-week-old control and and cKO mice. n=7/7 Control/KO. Data is presented as mean ± S.E.M. **d,** H&E staining of cross sections of descending aortas from control and a and cKO mice. Representative of n=3. Scale bar: 100 μm. A: tunica adventitia, M: tunica media, L: lumen. **e,** Average media thickness of descending aorta in control and cKO mice. N=3 mice per group, average of 6 equally spaced-out measurements per mouse aorta, using imageJ. Data is presented as mean ± S.D. **f,** Verhoeff Van Gieson stain for elastin, representative of n=3 control and and cKO mice. Scale bar: 100 μm. A: tunica adventitia, M: tunica media, L: lumen. **g,** Tail cuff blood pressure measurements of 10-week-old control and KO mice (n=9/7 control/KO). Each dot represents the average blood pressure of 5 days consecutive measurements per mouse. Data is represented as mean ± S.D. **h,** Heatmap of the top 200 upregulated and top 200 downregulated genes in control vs cKO (n=5 per group). Genes are colored according to their z-scores, with a selection of synthetic target genes and circulatory system development genes annotated. Gene ontology (GO) analysis of all downregulated genes in cKO vs control aortas is shown in the bottom right. Data represent n=5 per group. **i,** Expression MA plot as shown in Figure 2a, with unchanged SMC phenotypic transition genes annotated. **j,** UMAP visualization of gene expression in single cells in control and cKO aotae, showing SMC marker *Myh11, Prdm16* exon 9, Activated SMC cluster marker *Sod3,* pre-modulated SMC cluster marker *Vmp1,* and PRDM16 target genes *Col4a2, Chrdl1, Tnnt2, Sorbs1*. **k,** Immunostaining for MYH11 (magenta), PRDM16 (green), PECAM-1 (CD31, white), and DAPI (nuclei, blue), and in cross sections of descending aorta from control and and iKO mice. Representative of n=3/3 Ctrl/KO. Scale bar: 20 μm. M: tunica media, L: lumen. **l,** Tail cuff measurements of blood pressure in 10-week-old control and KO mice (n=8/7 control/KO). Each dot represents the average blood pressure of 4 days consecutive measurements per mouse. Data is presented as mean ± S.E.M. **m,** Average media thickness of descending aorta in control and iKO mice. N=3 mice per group, average of 6 equally spaced-out measurements per mouse aorta, using imageJ. Data is presented as mean ± S.D. **n,** H&E staining of cross sections of descending aortas from control and a and iKO mice. Representative of n=3. Scale bar: 200 μm. **o,** Verhoeff Van Gieson stain for elastin, representative of n=3 control and and iKO mice. Scale bar: 200 μm. A: tunica adventitia, M: tunica media, L: lumen. **p,** Heatmap of the top 200 upregulated and top 200 downregulated genes in control vs iKO (n=5 per group). Genes are colored according to their z-scores, with a selection of synthetic target genes and circulatory system development genes annotated. GO analysis of all downregulated genes in iKO vs control aortas is shown in the bottom right. Data represent n=3 per group. **q,** Expression MA plot as shown in Figure 2i, with unchanged SMC phenotypic transition genes annotated.

#### Supplemental Figure 3

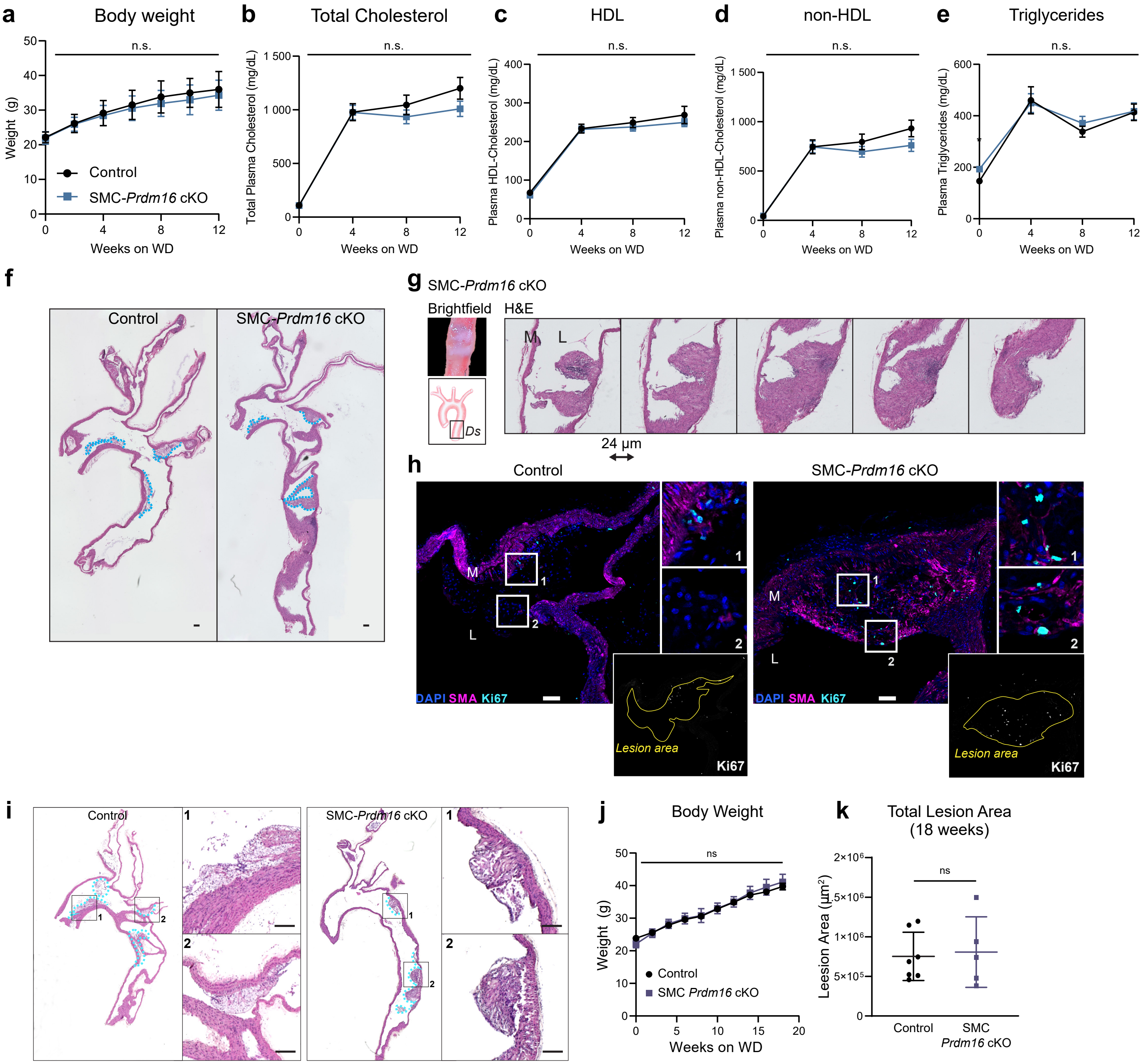


**Supplemental Figure 3 | a,** Bi-weekly body weight measurements of control and cKO mice from the time of AAV8-PCSK9-D377Y injection (8 weeks old) throughout the 12 weeks of western diet feeding (n = 15/16 Ctrl/KO). Data is represented as mean ± S.E.M. **b-e,** FPLC measurements of total cholesterol (a) high-density lipoprotein (HDL) (c), non-high density lipoprotein fraction (non-HDL) (d) and triglyceride (e) levels in 4h fasted plasma samples taken the day before PCSK9 injection (t=0) and throughout the 12 weeks of western diet feeding (n = 15/16 Ctrl/KO). Data is presented as mean ± S.E.M. **g,** H&E staining of sequential sagittal sections through an aortic lesion from and cKO mice. The brightfield image shows the gross shape of the lesion at dissection. M: tunica media, L: lumen. Verhoeff van Gieson staining for elastin in aortas from control and and cKO mice 12 weeks after inducing atherosclerosis. Representative of n = 3/3 Ctrl/KO. M: tunica media, L: lumen. **h,** Immunostaining for Ki67 (Cyan), smooth muscle actin/Acta2 (SMA, magenta), and Dapi (nuclei, Blue) in lesions from Control and KO mice. Representative of n=3 per group. Scale bar: 50 µm M: tunica media, L: lumen. **i,** Representative H&E staining of sagittal sections from the aortic arch of control and and cKO mice throughout 18 weeks of western diet feeding. Lesion area is outlined in blue. Only lesions in the arch, the start of the descending aorta and the beginnings of the branches were included. Scale bar: 100 µm. **i,** bi-weekly body weight measurements of control and cKO mice throughout 18 weeks of Western diet feeding. **k,** total lesion area of control and cKO mice (n = 7 / 5 respectively)

### Supplemental Figure 4

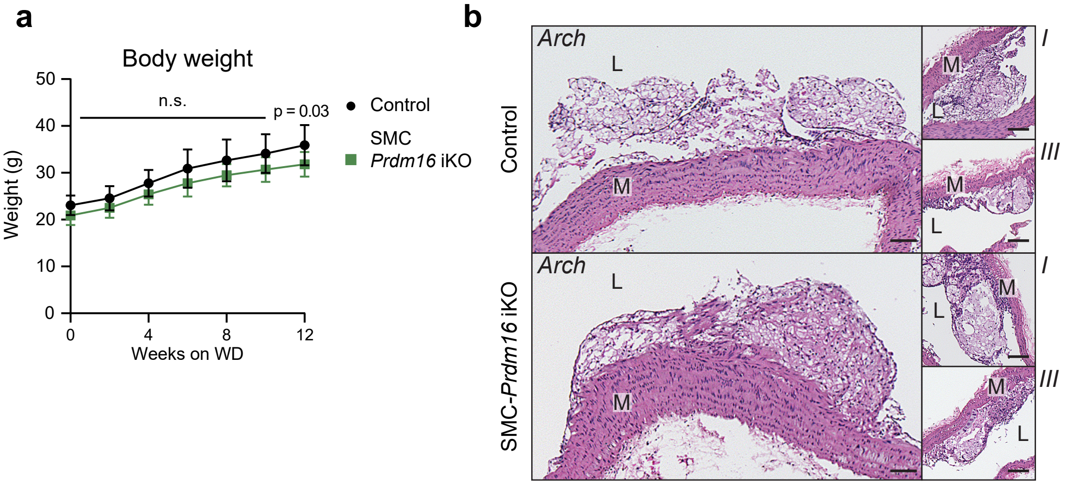


**Supplemental Figure 4 | a,** Body weight of control and KO mice from the time of PCSK9 injection (8 weeks old) throughout the 12 weeks of western diet feeding (n = 15/13 Ctrl/KO). Data is presented as mean ± S.E.M. ns = not significant. **b,** H&E-stained sagittal sections of the aortic arch from control and iKO mice containing atherosclerotic lesions at different locations. Images are representative of (n = 13/11 Ctrl/KO). Scale bar: 50 µm. A diagram of the aortic arch shows the orientation: I= brachiocephalic artery, III = left subclavian artery, Ds = descending aorta.

### Supplemental Figure 5

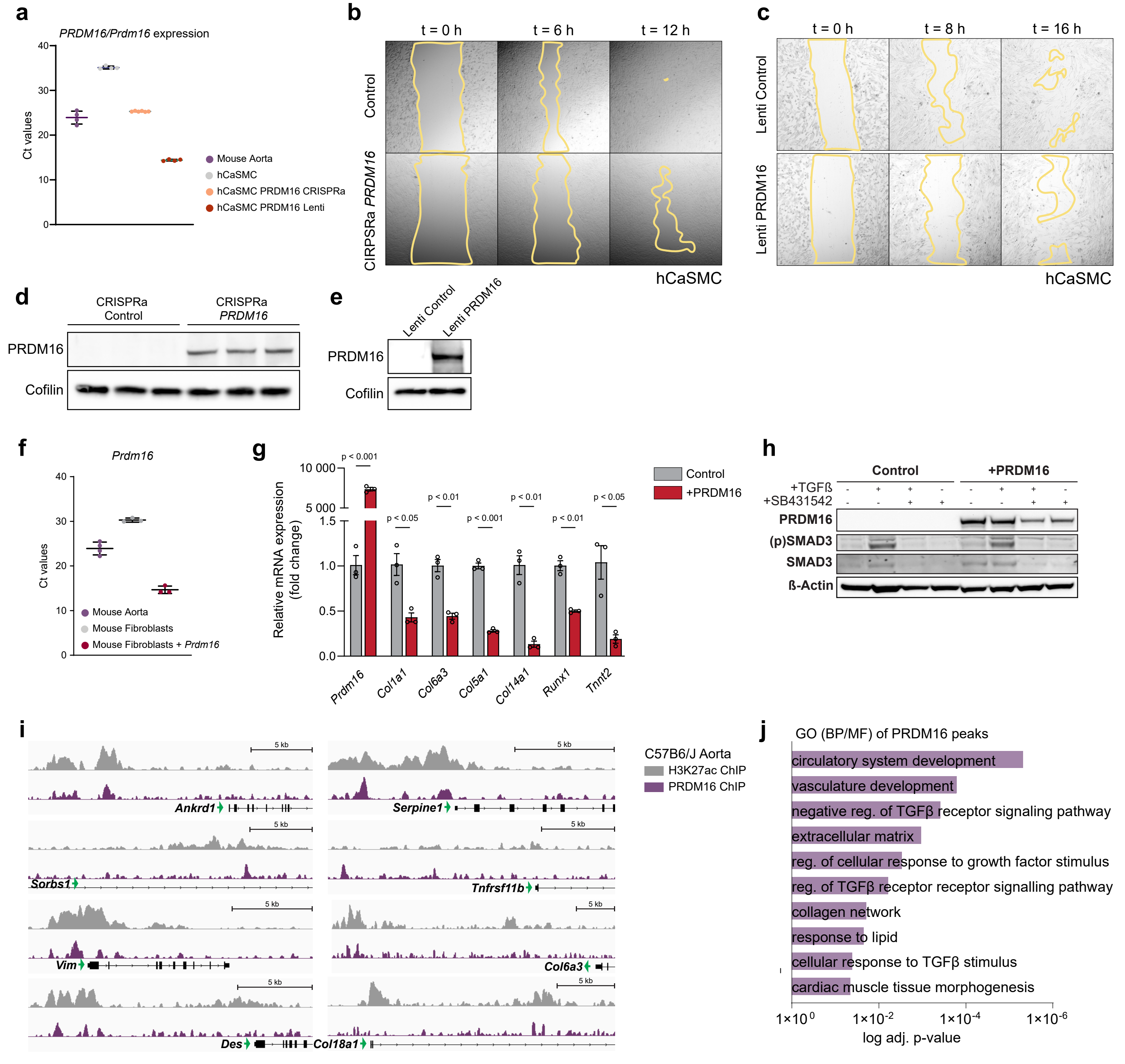


**k** Intersections for GO term names Prdm16 ChIP Peaks

| | | |
|---|---|---|
| circulatory system development | GO:0072359 | ECE1,DHRS3,COL4A1,PTGIS,SERPINE1,HEY2,COL8A1,PTPN14, SMAD7,SMAD6, AMOTL2,IL12B,NAGLU, RHOB,MYH11,HEXIM1,GJA4,ANKRD1, RRAS,EPN2,PECAM1,MYLK2, ERRFI1,NTRK3,SOCS3,COL18A1,CLIC4,EFNB2,COL5A1,HEYL,TRAF3IP2 |
| vasculature development | GO:0001944 | COL4A1,PTGIS,SERPINE1,HEY2,COL8A1,PTPN14,SMAD7,SMAD6, AMOTL2,IL12B,NAGLU,RHOB, GJA4,RRAS, EPN2,PECAM1, ERRFI1,SOCS3,COL18A1,CLIC4,EFNB2,COL5A1 |
| negative regulation of transforming growth actor beta receptor signaling pathway | GO:0030512 | PMEPA1,SMAD7,SMAD6,WFIKKN2,LDLRAD4,HTRA3,VASN |
| extracellular matrix | GO:0031012 | COL4A1,SERPINE1,NCAM1,COL8A1,MGP,CD180, CRISPLD2,VASN,THSD4,COL6A3,ANXA11, TNFRSF11B,COL18A1, MMP16,SOD3,COL5A1 |
| regulation of cellular response to growth factor stimulus | GO:0090287 | SULF2,PMEPA1,SMAD7,SMAD6,IL12B,WFIKKN2,LDLRAD4,HTRA3,EPN2,VASN |
| regulation of transforming growth factor beta receptor signaling pathway | GO:0017015 | PMEPA1,SMAD7,SMAD6,WFIKKN2,LDLRAD4,HTRA3,VASN |
| collagen network | GO:0098645 | COL4A1,COL8A1 |
| response to lipid | GO:0033993 | TPCN2,PMEPA1,SERPINE1,ADCY5,SMAD6,IRS1,IL12B,CD180,ARID5A, NAGLU,NR1D1,GBP5,ANKRD1,CCR7, PDK4,ERRFI1,DDIT4,INHBB,VIM,TRAF6 |
| ellular response to transforming growth factor beta stimulus | GO:0071560 | PMEPA1,SMAD7,SMAD6,WFIKKN2,LDLRAD4,HTRA3,ANKRD1,VASN |
| cardiac muscle tissue morphogenesis | GO:0055008 | HEY2,SMAD7,NAGLU,ANKRD1,MYLK2 |

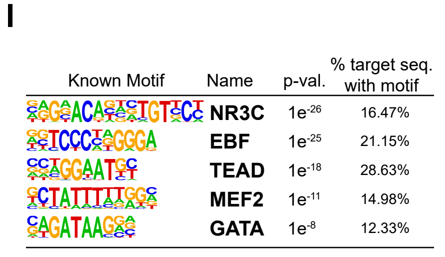

**Supplemental Figure 5 | a,** Ct values for *PRDM16/Prdm16* mRNA levels of equal RNA input in mouse aorta, hCaSMCs, as well as hCaSMCs transduced with *PRDM16* CRISPRa or Lenti-PRDM16. **b,** Widefield images of the scratch assay shown in Fig 6a. Images are stills of a movie at 0, 6, 12 h after infliction of scratch. The yellow lines mark the wound healing borders. **c,** Widefield images of the scratch assay shown in Fig 6d. Images are stills of a movie at 0, 6, 12 h after infliction of scratch. The yellow lines mark the wound healing borders. **c,** immunoblot showing PRDM16 protein levels in control hCaSMCs and hCaSMCs expressing a *PRDM16* CRISPRa construct. Cofilin is used as a loading control. N=3 **e,** immunoblot showing PRDM16 protein levels in control hCaSMCs and hCaSMCs expressing lenti-PRDM16. Representative of n=3. Cofilin is used as a loading control. **f,** Ct values for *Prdm16* mRNA levels of equal RNA input in mouse aorta, mouse fibroblasts and fibroblasts stably expressing *Prdm16*. **g,** mRNA levels of indicated genes in control and PRDM16-expressing fibroblasts. N = 3. **h,** Immunoblot for PRDM16, SMAD3 and phosphorylated SMAD3 ((p)SMAD3) in lysates from PRDM16-expressing and control cells treated with TGFß and/or ß-actin was used as a loading control. Image is representative of n = 3. **i,** ChIP tracks for PRDM16 and H3K27Ac in C57B6/J aortae at the indicated synthetic PRDM16 target genes. **g,** GO analysis of all genes containing a PRDM16 ChIP-seq peak. **h,** intersections of the GO analysis shown in (g). each gene name constitutes an identified ChIP-seq peak associated with the indicated GO terms. **i,** the top 5 most significantly enriched DNA motifs within the PRDM16 binding sites.
